# HORMA domain proteins and a Pch2-like ATPase regulate bacterial cGAS-like enzymes to mediate bacteriophage immunity

**DOI:** 10.1101/694695

**Authors:** Qiaozhen Ye, Rebecca K. Lau, Ian T. Mathews, Jeramie D. Watrous, Camillia S. Azimi, Mohit Jain, Kevin D. Corbett

## Abstract

Bacteria are continually challenged by foreign invaders including bacteriophages, and have evolved a variety of defenses against these invaders. Here, we describe the structural and biochemical mechanisms of a bacteriophage immunity pathway found in a broad array of bacteria, including pathogenic *E. coli* and *Pseudomonas aeruginosa*. This pathway employs eukaryotic-like HORMA domain proteins that recognize specific peptides, then bind and activate a cGAS/DncV-like nucleotidyltransferase (CD-NTase) to generate a cyclic tri-AMP (cAAA) second messenger; cAAA in turn activates an endonuclease effector, NucC. Signaling is attenuated by a homolog of the AAA+ ATPase Pch2/TRIP13, which binds and likely disassembles the active HORMA-CD-NTase complex. When expressed in non-pathogenic *E. coli*, this pathway confers immunity against bacteriophage λ infection. Our findings reveal the molecular mechanisms of a bacterial defense pathway integrating a cGAS-like nucleotidyltransferase with HORMA domain proteins for threat sensing through protein detection, and negative regulation by a Pch2-like ATPase.

## Introduction

All cellular life depends on signaling pathways that can rapidly sense and respond to changes in both a cell’s internal state and its external environment. Many such signaling pathways rely on diffusible second messenger molecules, which are synthesized by sensor proteins and activate effector proteins to drive a physiological response. In bacteria, the cyclic dinucleotide second messengers cyclic di-AMP, cyclic di-GMP, and cyclic GMP-AMP (cGAMP) play key roles in cellular homeostasis and pathogenesis (Corrigan and Gründling, 2013; Davies et al., 2012; Hengge, 2009). In mammals, cGAMP and linear oligoadenylate are important innate-immune signals, synthesized by cGAS (<u>c</u>yclic <u>G</u>MP-<u>A</u>MP <u>s</u>ynthase) and OAS (<u>o</u>ligoadenylate <u>s</u>ynthase) family proteins in response to cytoplasmic DNA or double-stranded RNA (Chen et al., 2016; Hornung et al., 2014). Nucleotide-based second messengers also mediate cross-kingdom signaling, with mammalian innate immune receptors able to recognize and respond to a variety of bacterially-generated second messengers (Burdette et al., 2011; McFarland et al., 2017; Whiteley et al., 2019).

A major family of second messenger biosynthetic enzymes found across kingdoms is distantly related to DNA polymerase β and includes bacterial DncV (<u>D</u>i<u>n</u>ucleotide *c*yclase in *<u>V</u>ibrio*), which synthesizes cGAMP (Davies et al., 2012; Zhu et al., 2014), and mammalian cGAS and OAS. Recently, a comparative genomics analysis identified hundreds of distinct cGAS/DncV-like nucleotidyltransferases (CD-NTases) in environmental and human patient-derived bacterial strains (Burroughs et al., 2015). A later biochemical survey showed that these enzymes synthesize a marked variety of second messengers, including cyclic dinucleotides with both purine and pyrimidine bases, as well as cyclic trinucleotides and other as-yet uncharacterized molecules (Whiteley et al., 2019). The majority of bacterial CD-NTases are found in operons associated with transposase or integrase genes and/or conjugation systems, and are sparsely distributed throughout environmental and pathogenic bacteria, suggesting that these operons are part of the “mobilome”. The presence of putative effector proteins in these operons including predicted nucleases, proteases, and phospholipases further suggests that the operons provide protection against foreign threats such as bacteriophages, plasmids, or other bacteria (Burroughs et al., 2015; Whiteley et al., 2019).

Biochemical analysis of 66 purified bacterial CD-NTases showed that the majority of these enzymes are inactive in vitro, suggesting that their activity is tightly regulated (Whiteley et al., 2019). Intriguingly, a subset of bacterial CD-NTase operons encode one or two proteins with weak homology to eukaryotic HORMA domain proteins, along with an ortholog of the AAA+ ATPase regulator of HORMA domain proteins, Pch2/TRIP13 (Burroughs et al., 2015). In eukaryotes, HORMA domains (named for the first three proteins shown to possess this domain: <u>Ho</u>p1, <u>R</u>ev7, and <u>Ma</u>d2) bind short peptides called “closure motifs” and assemble into signaling complexes that control inter-homolog recombination in meiosis (Hop1), DNA repair (Rev7), the mitotic spindle assembly checkpoint (Mad2), and autophagy (Atg13, Atg101) (Aravind and Koonin, 1998; Rosenberg and Corbett, 2015). The HORMA domain possesses two distinct folded states: an “open” state unable to bind closure motifs, and a “closed” state in which the HORMA domain C-terminus wraps entirely around a bound closure motif (Luo and Yu, 2008; Rosenberg and Corbett, 2015). The Pch2/TRIP13 ATPase disassembles HORMA:closure motif complexes by partially unfolding the N-terminus of the HORMA domain, dissociating the complex and in some cases converting the HORMA domain from the “closed” to the “open” state (Ma and Poon, 2016; Ye et al., 2015, 2017).

While HORMA domains play key signaling roles in eukaryotes, these proteins have not been identified in bacteria. The presence of putative HORMA domain proteins in bacterial signaling operons suggests that these operons may utilize the HORMA domain’s peptide binding and conformational conversion abilities to control second messenger synthesis by their cognate CD-NTases. Here, we show that this is indeed the case: bacterial HORMA domain proteins adopt the canonical HORMA domain fold, undergo conformational conversion between open and closed states, and bind specific closure motif peptides. Binding of a “closed” HORMA domain protein to its cognate CD-NTase activates synthesis of a cyclic tri-AMP (cAAA) second messenger, which activates an effector endonuclease also encoded in the operon. Bacterial Pch2-like ATPases function as negative regulators of signaling, binding and likely disassembling the active CD-NTase:HORMA complex. Finally, we show that a CD-NTase+HORMA+Pch2 operon encoded by a patient-derived *E. coli* strain confers immunity against bacteriophage λ infection. Together, these results define the molecular mechanisms of a new bacterial defense system that employs the first identified bacterial HORMA domain proteins as a novel regulator of second messenger signaling.

## Results

### Bacterial HORMA domain proteins bind and activate second messenger synthesis by their associated CD-NTases

Bacterial CD-NTases have been grouped into eight major clades denoted A-H, and further divided into numbered sub-clades, based on sequence similarity (Whiteley et al., 2019). CD-NTases in clades C01 and D05 are found in operons with one or two putative HORMA domain proteins, a Pch2/TRIP13-like ATPase, and one of several diverse putative effector proteins (Burroughs et al., 2015; Whiteley et al., 2019). We found that the putative HORMA domain proteins fall into three families: HORMA proteins from operons with a single putative HORMA protein (1-HORMA operons) form one family, while operons with two putative HORMA proteins (2-HORMA operons) contain one member of each of two diverged families (Fig. S1). Similarly, Pch2 proteins from 1-HORMA and 2-HORMA operons fall into two distinct families (Fig. S1). These data suggest that 1-HORMA and 2-HORMA operons, despite sharing similar components, have evolutionarily diverged from one another.

To address the function of bacterial HORMA domain protein-containing operons, we focused on two such operons: the 1-HORMA operon from *E. coli* strain MS115-1 whose CD-NTase falls into clade C01 (hereon termed CdnC), and the 2-HORMA operon from *Pseudomonas aeruginosa* strain ATCC27853 whose CD-NTase falls into clade D05 (CdnD) (Fig. 1A). The *E. coli* operon is found in several patient-derived *E. coli* strains, and the *P. aeruginosa* operon is found in over a quarter of patient-derived *P. aeruginosa* strains. Both operons also encode related putative endonuclease effectors, termed REase by Burroughs et al. (Burroughs et al., 2015) and here termed NucC (<u>Nuc</u>lease, <u>C</u>D-NTase associated). The *P. aeruginosa* ATCC27853 operon also encodes a putative structural maintenance of chromosomes (SMC)-family ATPase (not further examined here), and both operons have nearby toxin-antitoxin gene pairs (Fig. S2A and Fig. S2D).

**Figure 1.**
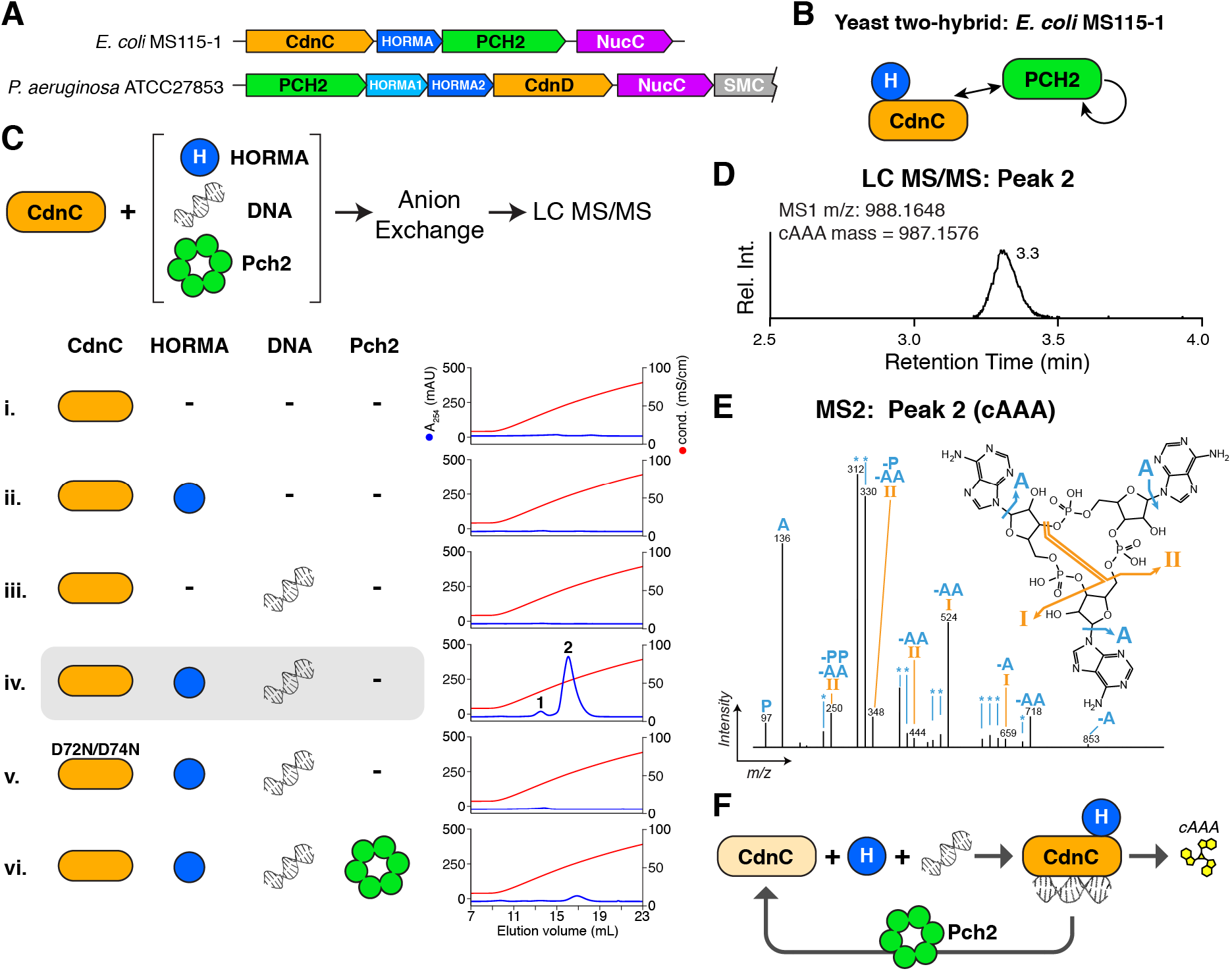
HORMA+Pch2-associated CD-NTases synthesize cAAA. **(A)** Schematic of CD-NTase+HORMA+Pch2 operons from *E. coli* MS115-1 (top) and P. *aeruginosa* ATCC27853 (bottom). See Fig. S1 for phylogenetic analysis of Pch2 and HORMA proteins. *P. aeruginosa* ATCC27853 encodes a Rad50-like SMC-family protein (SMC) not further studied here. See Table S2 for protein sequences. **(C)** Summary of yeast two-hybrid and three-hybrid assays with the *E. coli* MS115-1 CdnC, HORMA, and Pch2. See Fig. S2A for yeast two-hybrid and three-hybrid results, Fig. S2B for purification and stoichiometry of a CdnC:HORMA complex, and Fig. S2C for purification of a CdnC:HORMA:Pch2^EQ^ complex. See Fig. S2D-G for equivalent assays with the *P. aeruginosa* ATCC27853 operon. **(C)** *Top:* Schematic of second messenger synthesis assays. *Ec* CdnC was incubated with potential regulators (HORMA, plasmid DNA, and Pch2) plus ATP, then the products were separated by anion-exchange chromatography and analyzed by liquid chromatography with tandem mass spectrometry (LC-MS/MS). *Bottom:* Anion exchange elution profiles from second messenger synthesis assays using *Ec* CdnC, HORMA, DNA, and Pch2. Blue lines show absorbance at 254 nm, and red lines show conductivity for the 0.2-2.0 M ammonium acetate gradient elution. CdnC D72N/D74N contains aspartate-to-asparagine mutations in the putative active-site residues 72 and 74. See Fig. S3A for assays with *Ec* CdnC and different DNAs, and Fig. S3B for equivalent assays with *Pa* CdnD. **(D)** Liquid chromatography elution profile of the major product of *Ec* CdnC (peak 2 from panel C, sample iv), with measured MS1 m/z and theoretical mass of cAAA. **(E)** MS2 fragmentation spectrum of the major product of *Ec* CdnC, annotated according to expected fragments of cAAA. The m/z of the extracted ion (corresponding to the [M+H] adduct of cAAA was 988.1648, consistent with cAAA (monoisotopic mass = 987.1576 amu; molecular weight = 987.6263 amu). The minor product of *Ec* CdnC (peak 1 from panel C, sample iv) was also analyzed by LC MS/MS and confirmed to be cAA (not shown). **(F)** Model for CdnC activation: Binding of HORMA and DNA to CdnC activates cAAA production, and Pch2-mediated disassembly inactivates signaling.

As a first step, we performed yeast two-hybrid analysis with the *E. coli* MS115-1 (*Ec*) proteins, and found that *Ec* CdnC and *Ec* HORMA bind one another (Fig. 1B and Fig. S2A). In a yeast three-hybrid experiment, a mutant of *Ec* Pch2 that stabilizes hexamer formation and disrupts ATP hydrolysis (Walker B motif residue Glu159 to Gln; Pch2^EQ^) binds to an *Ec* CdnC:HORMA complex (Fig. 1B and Fig. S2A). Similar yeast two-hybrid analysis with the *P. aeruginosa* ATCC27853 *(Pa)* proteins showed that *Pa* HORMA1 and HORMA2 each interact with *Pa* CdnD, and also interact with one another (Fig. S2D). We also find that both *Ec* NucC and *Pa* NucC self-interact in yeast two-hybrid assays, suggesting that NucC forms homodimers or larger oligomers (Fig. S2A and Fig. S2D).

Both *Ec* CdnC and *Pa* CdnD were previously found to be inactive in vitro (Whiteley et al., 2019), leading us to test if their activity depends on their cognate HORMA and Pch2 proteins. For this purpose, we tested purified proteins in an in vitro second messenger synthesis assay based on prior work with a related CD-NTase, *E. cloacae* CdnD02 (Fig. 1C) (Whiteley et al., 2019). We first tested for *Ec* CdnC activity in the presence of *Ec* HORMA and *Ec* Pch2. As the related cGAS and OAS enzymes both require nucleic acid binding for activation (Hornung et al., 2014), we also included double-stranded DNA as a potential regulator. We find that while CdnC is inactive on its own (Fig. 1C, sample i), the enzyme synthesizes two phosphatase-resistant products in the presence of both *Ec* HORMA and DNA (Fig. 1C, sample iv). Addition of either *Ec* HORMA or DNA alone did not activate *Ec* CdnC, suggesting that the enzyme requires both activators. CdnC+HORMA was activated by long doublestranded DNAs, but not by single-stranded DNA or short (40 bp) doublestranded DNA (Fig. S3A). CdnC activity was disrupted by mutation of the putative active site residues Asp72 and Asp74, demonstrating that CdnC uses a similar biosynthetic mechanism as related CD-NTases (Fig. 1C, sample v). Finally, we found that addition of sub-stoichiometric amounts of *Ec* Pch2 (1:60 molar ratio of active Pch2 hexamers to CdnC+HORMA) strongly reduced second messenger synthesis (Fig. 1C, sample vi). Together, these data support a model in which *Ec* CdnC is activated by binding its cognate HORMA protein and DNA, and inactivated by Pch2, potentially through direct disassembly of a CdnC:HORMA complex (Fig. 1F).

We next purified the major and minor products of *Ec* CdnC and analyzed them by liquid chromatography coupled to tandem mass spectrometry (LC MS/MS). We found that the major product of *Ec* CdnC is cyclic tri-AMP (cAAA; Fig. 1D-E), and that the minor product is cyclic di-AMP (cAA; not shown). *Ec* CdnC synthesizes only cAAA/cAA even in the presence of both GTP and ATP, indicating that the enzyme uses only ATP as a substrate (not shown). Thus, *Ec* CdnC synthesizes a cyclic trinucleotide second messenger only when activated by its cognate HORMA domain protein and double-stranded DNA.

We next tested the activity of *Pa* CdnD in the presence of *Pa* HORMA1, *Pa* HORMA2, and *Pa* Pch2. While *Pa* CdnD shows lower activity than *Ec* CdnC in vitro and requires Mn^2+^ instead of Mg^2+^ as a cofactor, we find that CdnD also primarily synthesizes cAAA (Fig. S3B). *Pa* CdnD requires HORMA2 for activity, but is not activated by HORMA1 (Fig. S3B), perhaps reflecting functional specialization of the two HORMA domain proteins in this operon. As with the *E. coli* proteins, addition of *Pa* Pch2 to a reaction with *Pa* CdnD and HORMA2 strongly inhibits second messenger synthesis (Fig. S3B). Finally, we find that in contrast to *Ec* CdnC, DNA is not required for *Pa* CdnD activity, nor does addition of DNA further stimulate cAAA production by the enzyme (Fig. S3B). Thus, while *Ec* CdnC and *Pa* CdnD are regulated differently, the two enzymes share a common dependence on HORMA domain protein binding for activation, and generate the same cyclic trinucleotide second messenger.

### Structures of HORMA-associated CD-NTases

To address the structural and biochemical mechanisms by which *Ec* CdnC and *Pa* CdnD synthesize cAAA in a regulated manner, we first purified and determined high-resolution crystal structures of both enzymes (Fig. 2A-B). Both *Ec* CdnC and *Pa* CdnD adopt a polymerase β-like nucleotidyltransferase fold, with two lobes sandwiching a large active site. *Ec* CdnC and *Pa* CdnD adopt similar overall structures, with a 3.95 Å Cα r.m.s.d. (over 250 residues), and are more structurally related to one another than to either DncV (clade A) or CdnE, both of which synthesize cyclic dinucleotide second messengers (Fig. 2C).

**Figure 2.**
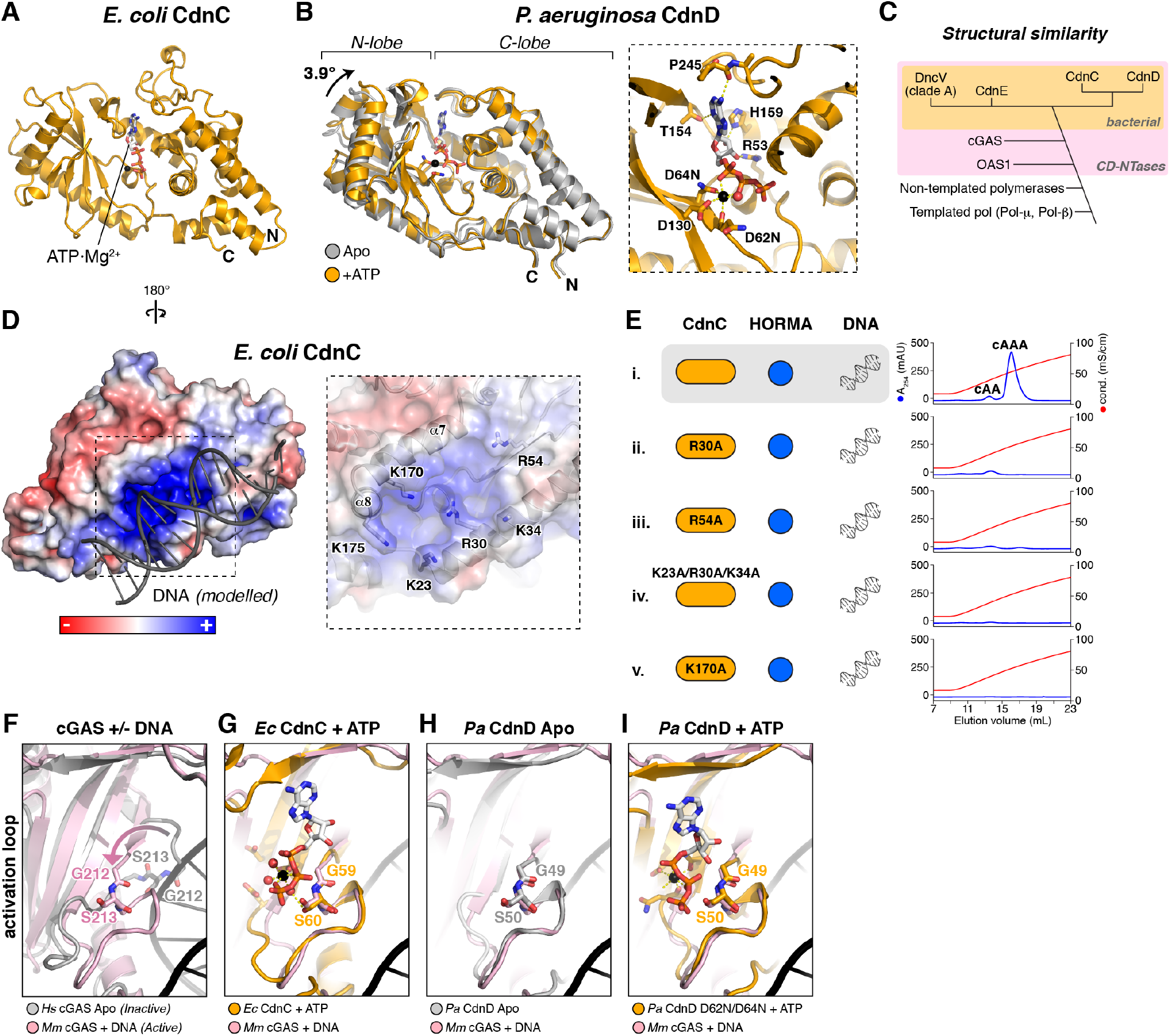
Structures of HORMA+Pch2-associated CD-NTases. **(A)** Structure of *Ec* CdnC, with bound ATP·Mg^2+^ shown as sticks. **(B)** Overlay of *Pa* CdnD in the Apo state (gray) and bound to ATP (orange), with bound ATP·Mg^2+^ shown as sticks. *Right:* Closeup view of ATP·Mg^2+^ binding to *Pa* CdnD. See Fig. S4A for ITC assays measuring nucleotide binding to *Pa* CdnD. **(C)** Schematic of structural similarity in Pol-β type nucleotidyltransferases. CD-NTases are shaded in pink, and bacterial CD-NTases are shaded in orange. **(D)** Reverse view of *Ec* CdnC, showing surface charge and DNA (gray) modelled from a superposition with *M. musculus* cGAS bound to DNA (PDB 4K9B) (Gao et al., 2013). For surface conservation of *Ec* CdnC, see Fig. S4B. For surface conservation and charge of *Pa* CdnD, see Fig. S4C-D. Closeup view (right) shows positively-charged residues, with helices α7 and α8 (equivalent to mammalian cGAS DNA-binding surface; see Fig. S4E) labeled. **(E)** Anion exchange elution profiles from second messenger synthesis assays with wild-type *Ec* CdnC (sample i) and mutants to the putative DNA-binding surface (samples ii-v). **(F)** Overlay of inactive *H. sapiens* cGAS (gray; PDB ID 4O69 (Zhang et al., 2014)) with active, DNA-bound *M. musculus* cGAS (pink; PDB ID 4O6A (Zhang et al., 2014)) showing motion of the cGAS activation loop upon DNA binding. **(G)** Overlay of ATP-bound *Ec* CdnC (orange) with active cGAS (pink). **(H)** Overlay of Apo *Pa* CdnD (gray) with active cGAS (pink). **(I)** Overlay of ATP-bound *Pa* CdnD (orange) with active cGAS (pink).

In *Ec* CdnC, the active site is occupied by one molecule of ATP that was co-purified with the protein (Fig. 2A); we were unable to remove this tightly-bound nucleotide. We could purify *Pa* CdnD without a bound nucleotide, and found by isothermal titration calorimetry that the enzyme binds strongly to ATP, but not to GTP (Fig. S4A). These data agree with our findings that both *Ec* CdnC and *Pa* CdnD primarily synthesize cAAA. We determined structures of *Pa* CdnD in both Apo and ATP-bound states, and found that the two states differ by a 3.9° rotation of the enzyme’s N- and C-lobes with respect one another, with the ATP-bound state more “closed” than the Apo state (Fig. 2B).

Our biochemical data show that *Ec* CdnC requires dsDNA for activation, similar to mammalian cGAS (Fig. 1C). We modelled DNA binding to CdnC by overlaying our structure with a cGAS:DNA complex (Gao et al., 2013). In the resulting model, DNA contacts a surface of *Ec* CdnC that is strongly positively charged and highly conserved (Fig. 2D and Fig. S4B). To test the idea that this surface is responsible for DNA binding, we mutated several positively-charged residues on this surface of CdnC and measured cAAA synthesis. We found that mutations to the putative DNA binding surface eliminated cAAA production by CdnC in the presence of HORMA+DNA, supporting a role for this surface in DNA binding and activation of CdnC (Fig. 2E).

In mammalian cGAS, DNA binding to α-helices α7 and α8 restructures an “activation loop” in the enzyme’s active site to promote substrate binding and second messenger synthesis (Fig. 2F) (Civril et al., 2013; Kato et al., 2013; Li et al., 2013; Zhang et al., 2014). In our structures of *Ec* CdnC and *Pa* CdnD, the conformation of the activation loop is equivalent to the activated state of cGAS (Fig. 2G-I). Thus, while DNA binding is required for activation of *Ec* CdnC, the structural mechanism of activation is likely distinct from that of mammalian cGAS.

### Bacterial HORMA domain proteins adopt canonical open and closed states

We next sought to determine whether the putative bacterial HORMA domain proteins share the canonical HORMA domain fold and mechanism of closure motif binding (Fig. 3A). We first determined a structure of *Pa* HORMA2, and found that the protein adopts a canonical “closed” HORMA domain structure, with a central three-stranded β-sheet (β-strands 4, 5, and 6) backed by three α-helices (αA, B, and C) (Fig. 3B). Characteristic of closed HORMA domains, the N-terminus of *Pa* HORMA2 is disordered, and the C-terminus wraps around the protein and forms a β-hairpin (β-strands 8′ and 8″) against strand β5. Strikingly, the C-terminal safety belt of *Pa* HORMA2 embraces a peptide from the disordered N-terminus of a crystallographic symmetry-related monomer in the canonical closure motif binding site (Fig. 3B). Thus, *Pa* HORMA2 represents a *bona fide* HORMA domain with a closure-motif binding C-terminal safety belt.

**Figure 3.**
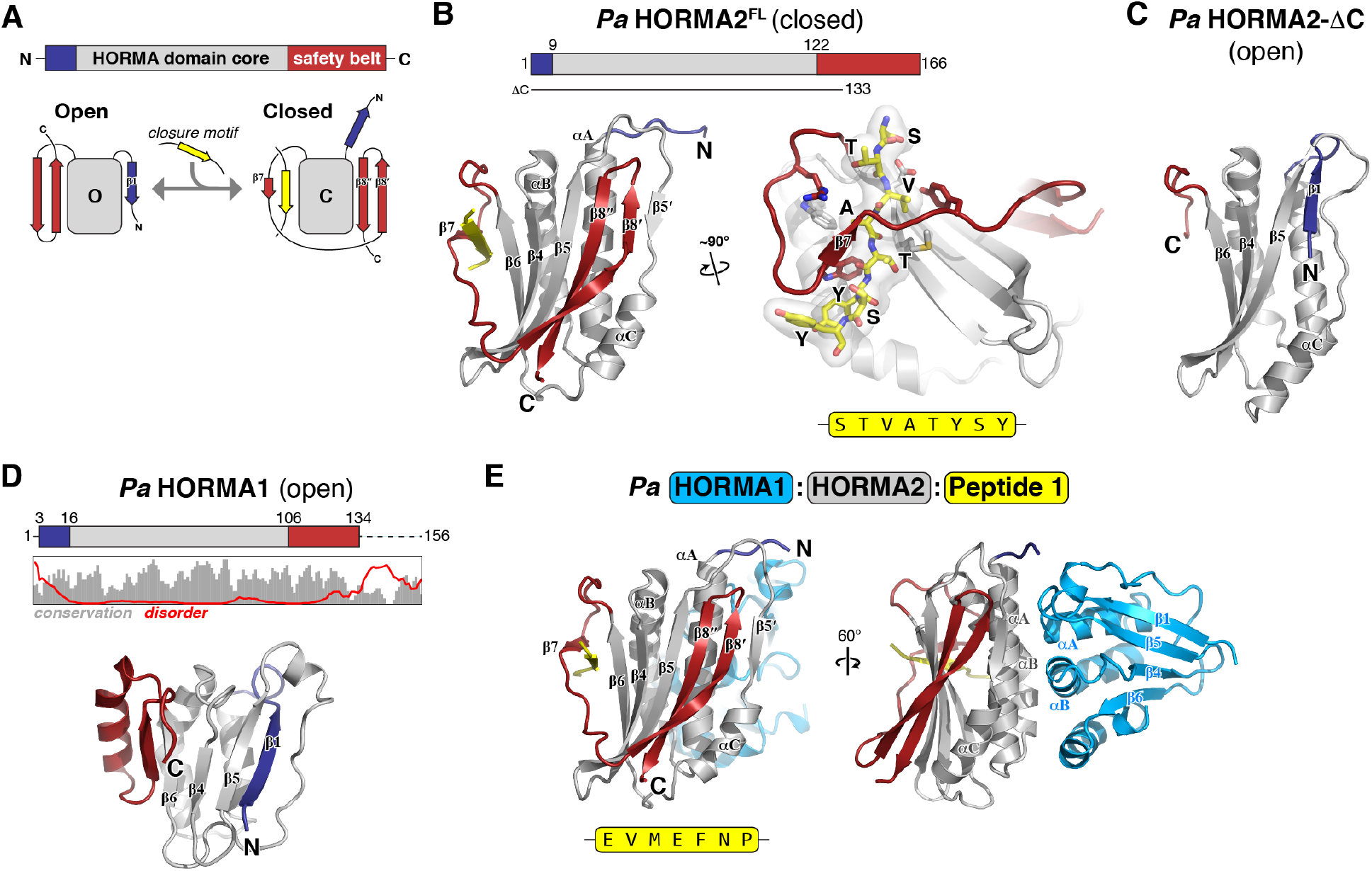
Structure and closure motif binding of bacterial HORMA proteins. **(A)** *Top:* Schematic of HORMA domain primary structure, with the HORMA domain core gray, N-terminus blue, and C-terminal safety belt red. *Bottom:* Schematic of the open and closed states of the canonical HORMA domain protein MAD2. The transition from open to closed involves movement of both the N-terminus (blue) and C-terminal safety belt (red), and binding of a closure motif peptide (yellow) by the safety belt. **(B)** *Left:* Structure of *Pa* HORMA2 in the closed state, colored as in panel (A). Secondary-structure elements are labeled as in MAD2, except for strand β5’, which is not observed in other HORMA domain proteins. *Right:* Closeup view of *Pa* HORMA2 binding the extended N-terminus of a symmetry-related *Pa* HORMA2 molecule, which mimics a closure motif (yellow). **(C)** Structure of *Pa* HORMA2-ΔC (lacking residues 134-166) in the open state. **(D)** *Top:* Schematic of *Pa* HORMA1, with plot showing the Jalview alignment conservation score (3-point smoothed; gray) (Livingstone and Barton, 1993) and DISOPRED3 disorder propensity (red) for aligned bacterial HORMA1 proteins (Jones and Cozzetto, 2015). *Bottom:* Structure of *Pa* HORMA1 in the open state. **(E)** Structure of the *Pa* HORMA1:HORMA2:Peptide 1 complex. See Fig. S5 for identification of closure motif sequences for *Pa* HORMA2 and *Ec* HORMA by phage display.

To identify preferred closure motif sequences for bacterial HORMA proteins, we performed phage display with purified *Pa* HORMA1, *Pa* HORMA2, and *Ec* HORMA and a library of 7-mer peptides. We identified consensus binding motifs enriched in hydrophobic amino acids for both *Pa* HORMA2 and *Ec* HORMA (Fig. S5A-B), and confirmed binding of *Pa* HORMA2 to a consensus peptide (Peptide 1: EVMEFNP; Fig. S5C). We have so far been unable to purify a stable complex of *Ec* HORMA with a closure motif peptide, suggesting it may bind closure motifs more weakly than *Pa* HORMA2. Nonetheless, the identification of a preferred binding sequence for *Ec* HORMA lends strong support to the idea that this protein, like *Pa* HORMA2, can adopt the closed state and bind closure motif peptides.

To determine whether *Pa* HORMA2 can also adopt an “open” HORMA domain state, we generated a truncated construct lacking the C-terminal safety belt residues 134-166 *(Pa* HORMA2-ΔC). The crystal structure of *Pa* HORMA2-ΔC shows that in contrast to full-length *Pa* HORMA2, the N-terminus of *Pa* HORMA2-ΔC is well-ordered and packs as a β-strand (β1) against strand β5 (Fig. 3C and Fig. S5D). This transition of the N-terminus from disordered to an ordered β1 strand is characteristic of open HORMA domains (Rosenberg and Corbett, 2015). Thus, bacterial HORMA domain proteins can adopt both the open and closed conformation, and bind closure motif peptides equivalently to their eukaryotic relatives.

We next determined a structure of *Pa* HORMA1, and found that this protein also adopts an open state with an ordered N-terminal β1 strand (Fig. 3D). Based on the sequence of the *Pa* HORMA1 N-terminus and the predicted disorder of its poorly-conserved C-terminus (Fig. 3D), we propose that this protein may be “locked” in the open state similarly to the eukaryotic HORMA domain protein Atg101 (Qi et al., 2015; Suzuki et al., 2015). Consistent with this idea, we were unable to identify *Pa* HORMA1-binding peptide sequences by phage display (not shown). Our data therefore shows that for 2-HORMA bacterial operons, one HORMA protein (e.g. *Pa* HORMA2) can adopt the open and closed states, bind closure motif peptides, and activate its cognate CD-NTase, while the second HORMA protein (e.g. *Pa* HORMA1) adopts an open state and does not activate its cognate CD-NTase.

We next determined a structure of a *Pa* HORMA1:HORMA2:Peptide 1 complex. Strikingly, the *Pa* HORMA1:HORMA2 dimer bears no resemblance to dimers of eukaryotic HORMA domains, which assemble through their aC helices (Mapelli et al., 2007; Rosenberg and Corbett, 2015). Rather, *Pa* HORMA1 and *Pa* HORMA2 interact through both proteins’ αA and αB helices, plus the short α-helix in the C-terminal region of *Pa* HORMA1 (Fig. 3E). The interface does not involve any elements of *Pa* HORMA2 that change conformation significantly between the open and closed states, and yeast two-hybrid analysis shows that the open *Pa* HORMA2-ΔC construct can also bind *Pa* HORMA1 (Fig. S5E-F). Thus, while *Pa* HORMA1 apparently does not bind closure motif peptides or undergo conformational conversion, it can likely scaffold larger complexes by binding *Pa* HORMA2 and CdnD.

### HORMA proteins bind their cognate CD-NTases specifically in the closed state

We next sought to understand how bacterial HORMA domain proteins regulate their cognate CD-NTases. We first purified and determined the structure of a complex containing *Pa* CdnD, HORMA2, and Peptide 1, and found that *Pa* HORMA2 binds the C-lobe of CdnD (Fig. 4A). Compared to its Apo state, HORMA2-bound *Pa* CdnD shows a 6.6° rotation of the N-lobe toward the C-lobe, consistent with the idea that binding of HORMA2 may aid activation of CdnD. HORMA2 interacts with CdnD through its β8′ strand and αC helix (Fig. 4B); as the β8′ strand does not form in the open HORMA domain state, this binding mode is likely dependent on HORMA2 adopting a closed conformation. Indeed, we found by yeast two-hybrid analysis that *Pa* HORMA2-ΔC, which adopts an open conformation, does not interact with *Pa* CdnD (Fig. 4C). Consistent with our ability to reconstitute a CdnD:HORMA1:HORMA2 complex, we could model a structure of *Pa* HORMA2 bound to both CdnD and HORMA1 with no apparent clashes (Fig. S6A).

**Figure 4.**
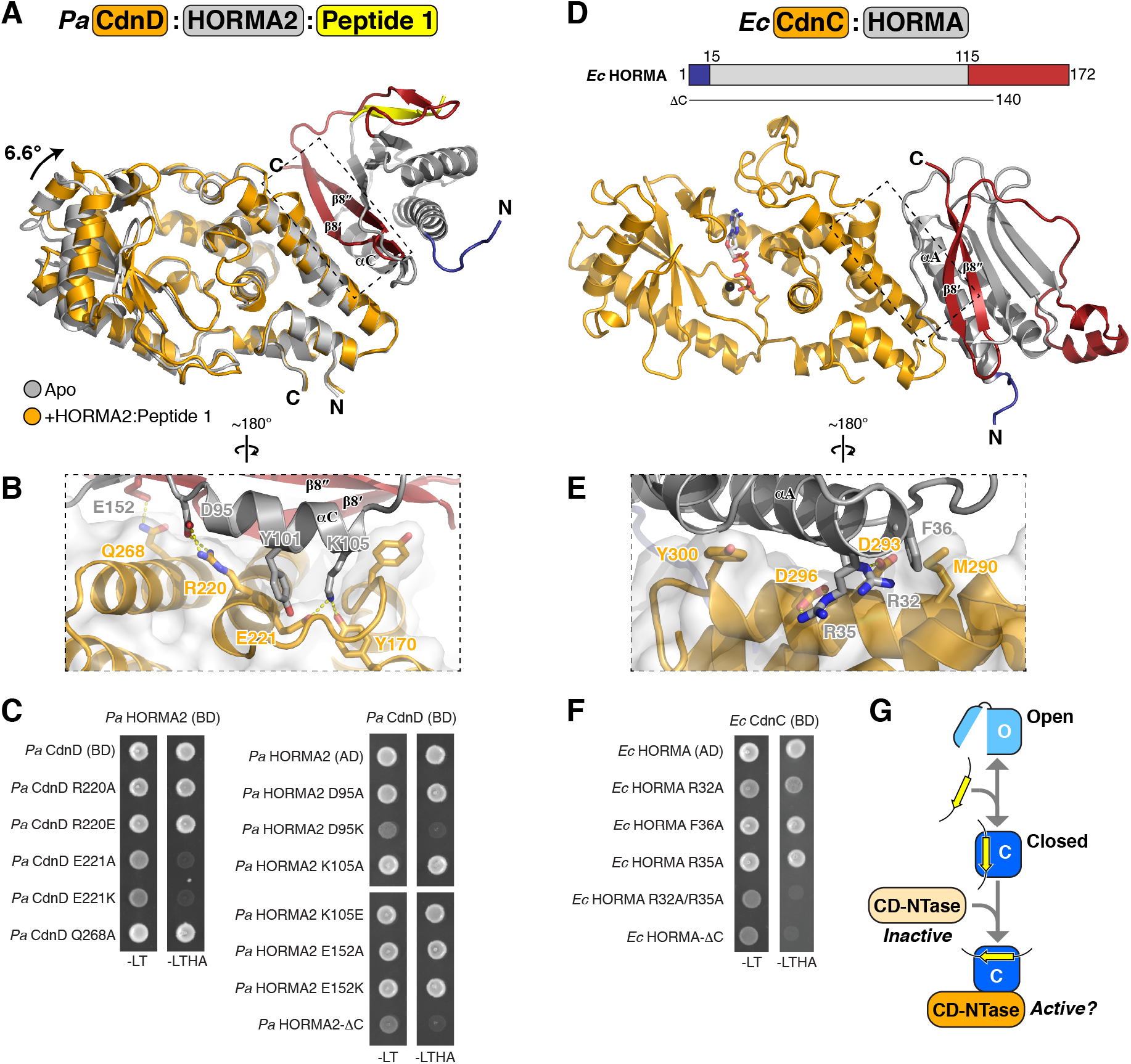
Structures of CD-NTase:HORMA complexes. **(A)** Overall structure of the *Pa* CdnD:HORMA2:Peptide 1 complex, with CdnD orange, HORMA2 gray with safety-belt red and N-terminus blue, and Peptide 1 yellow. CdnD is overlaid with Apo CdnD (gray), showing the 6.6° N-lobe rotation in the HORMA2-bound state. **(B)** Closeup view of the *Pa* CdnD:HORMA2 interface. **(C)** Yeast two-hybrid analysis of the *Pa* CdnD-HORMA2 interaction. BD: fusion to the Gal4 DNA-binding domain; AD: fusion to the Gal4 activation domain. -LT: media lacking leucine and tryptophan (non-selective); -LTHA: media lacking leucine, tryptophan, histidine, and adenine (stringent selection). **(D)** Overall structure of the *Ec* CdnC:HORMA complex, from a CdnC:HORMA:Pch2^EQ^ complex structure, with CdnC orange and HORMA gray with safety-belt red and N-terminus blue. **(E)** Closeup view of the Ec CdnC:HORMA interface. **(F)** Yeast two-hybrid analysis of the *Ec* CdnC:HORMA interaction. BD: fusion to the Gal4 DNA-binding domain; AD: fusion to the Gal4 activation domain. -LT: media lacking leucine and tryptophan (non-selective); -LTHA: media lacking leucine, tryptophan, histidine, and adenine (stringent selection). **(G)** Schematic of proposed CD-NTase activation. A soluble HORMA protein in the open state (light blue) binds a closure motif peptide (yellow), forming a closed-state complex (dark blue) that binds and activates CD-NTase.

We were unable to crystallize the isolated *Ec* CdnC:HORMA complex, but succeeded in crystallizing and determining a 2.6 Å resolution structure of an *Ec* CdnC:HORMA:Pch2^EQ^ complex (Pch2^EQ^ interactions are described below). In this three-protein complex, the structure of *Ec* HORMA closely resembles *Pa* HORMA2 in the closed state, with a disordered N-terminus and a well-ordered C-terminal safety belt forming a β-hairpin (β8′-8″) against strand β5 (Fig. 4D and Fig. S6B). Notably, in this structure *Ec* HORMA adopts an “empty” closed state, without a bound closure motif peptide. We modelled binding of a consensus closure motif sequence (HAAILFT) to *Ec* HORMA based on the *Pa* HORMA2:Peptide 1 structure and found that the four hydrophobic residues in the *Ec* HORMA-binding consensus would all fit into hydrophobic pockets on *Ec* HORMA (Fig. S6B), supporting the idea that the phage display experiment identified sequences that bind *Ec* HORMA as a closure motif.

The interaction of *Ec* HORMA with *Ec* CdnC closely resembles that of *Pa* HORMA2 with *Pa* CdnD (Fig. 4D). *Ec* HORMA binds the CdnC C-lobe, adjacent to CdnC’s putative DNA-binding surface. *Ec* HORMA binds CdnC through a similar interface as the one used by *Pa* HORMA2, binding CdnC largely through its αA helix (Fig. 4D-E). *Ec* HORMA’s β8’ strand does not contact CdnC, yet truncation of the protein’s C-terminus (*Ec* HORMA-ΔC, lacking residues 141-172) disrupts CdnC binding, suggesting that the *Ec* HORMA-CdnC interaction likely also depends on the HORMA protein adopting the closed state (Fig. 4F). Overall, our structures of *Pa* CdnD:HORMA2 and *Ec* CdnC:HORMA support a mechanistic model reminiscent of eukaryotic HORMA domain proteins, where conformational conversion of soluble open-state HORMA domain proteins to the closed state (e.g. upon closure motif peptide binding) enables binding to, and activation of, each HORMA domain protein’s cognate CD-NTase (Fig. 4G).

### Structure of a bacterial Pch2-like ATPase engaged with its CD-NTase:HORMA substrate

The Pch2-like ATPase encoded by bacterial HORMA-containing operons negatively regulates the production of cAAA by the HORMA-CD-NTase complex (Fig. 1C). To address the structural mechanism of this activity, and to compare this mechanism to that of eukaryotic Pch2/TRIP3 in control of the eukaryotic HORMA domain protein Mad2, we determined the structure of an *Ec* CdnC:HORMA:Pch2^EQ^ complex (Fig. 5A; the Pch2 E159Q mutation stabilizes hexamer assembly and substrate interactions by disallowing ATP hydrolysis (Ye et al., 2015, 2017)). The crystal structure of *Ec* CdnC:HORMA:Pch2^EQ^ reveals a mode of substrate binding and engagement for bacterial Pch2 that is similar to eukaryotic Pch2/TRIP13, with the high resolution of 2.6 Å providing unprecedented insight into substrate engagement and unfolding by AAA+ ATPases.

**Figure 5.**
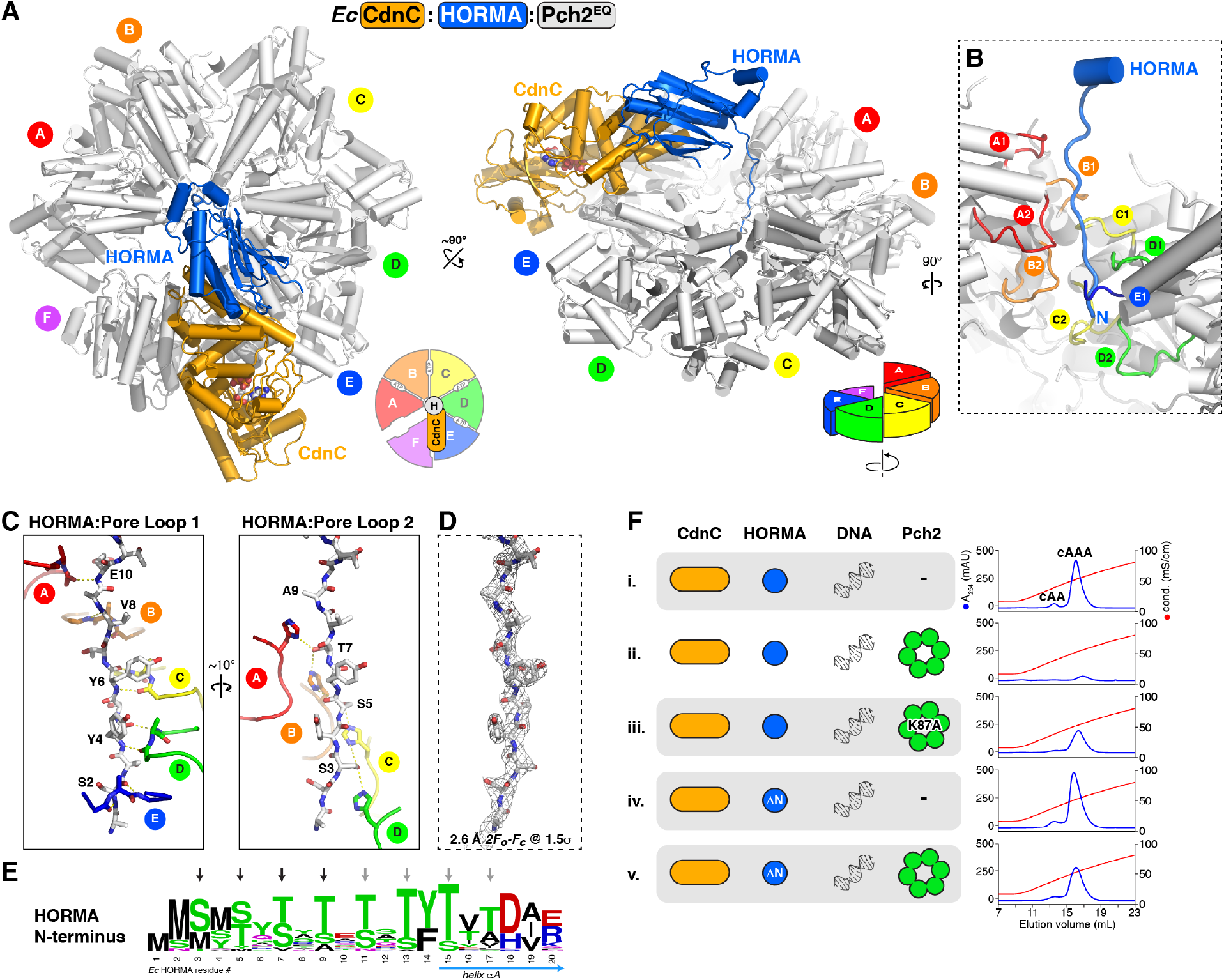
Structure of the *Ec* CdnC:HORMA:Pch2^EQ^ complex. **(A)** Two views (top view at left, side view at right) of the *Ec* CdnC:HORMA:Pch2^EQ^ complex, with CdnC orange, HORMA blue, and Pch2^EQ^ white. See Fig. S8 for views of the Pch2^EQ^ hexamer, conformational differences between subunits within the hexamer, and ATP binding. Pch2 forms a spiral conformation, with its A subunit at top and F subunit at bottom (see schematics). CdnC binds the top surface of Pch2 subunits E and F (see Fig. S8B for buried surface area). **(B)** Closeup view of the *Ec* HORMA N-terminus (blue) engaged by Pch2 pore loops 1 and 2 from chains A (red), B (orange), C (yellow), D (green, and E (blue; pore loop 1 only). **(C)** Detail views of interactions between the *Ec* HORMA N-terminus (white sticks) with Pch2 pore loop 1 (left) and pore loop 2 (right). HORMA residues Ser4, Tyr4, Tyr6, Val8, and Glu10 form an extended backbone hydrogen-bonding network with Pch2 pore loop 1 residues Gly119 (main-chain carbonyl) and Val121 (main-chain amine) from monomers A-E (Pch2 Arg120 side-chains are not shown for clarity). HORMA residues Ser3, Ser5, Thr7, and Ala9 form an extended hydrogen-bond network with Pch2 pore loop 2 residue His173 from monomers A-D. **(D)** View equivalent to panel (C) showing *2Fo-Fc* electron density for the HORMA N-terminus at 1.5 σ (2.6 Å resolution). **(E)** Sequence logo showing conservation of the N-termini of HORMA proteins from 1-HORMA operons. See Fig. S1C-D for analysis of Pch2 pore loop 1 and 2 conservation, and equivalent logos of Pch2 and HORMA from two-HORMA operons. **(F)** Anion exchange elution profiles from second messenger synthesis assays with wild-type proteins (samples i-ii), Pch2 Walker A mutant K87A (sample iii), and HORMA-ΔN (missing N-terminal residues 1-12; samples iv-v). and mutants to the putative DNA-binding surface (samples ii-v). See Fig. S3B for equivalent assays with *Pa* CdnD and Pch2 mutants.

A number of recent studies have reported moderate-resolution (3.2-5.0 Å) cryo-electron microscopy structures of AAA+ ATPase remodelers engaged with their substrates, including mammalian TRIP13 bound to a p31^comet^:Mad2 complex (Alfieri et al., 2018; Deville et al., 2017; Gates et al., 2017; Han et al., 2017; Puchades et al., 2017; Ripstein et al., 2017; White et al., 2018; Yu et al., 2018). These structures have revealed that active AAA+ remodelers adopt a right-handed spiral or “lock-washer” conformation, with 4-5 nucleotide-bound subunits forming a tight helical spiral and engaging an extended substrate peptide with their “pore loops”. In the case of TRIP13, the structures revealed that the “adapter” protein p31^comet^ binds the top surface of the TRIP13 hexamer, thereby positioning the substrate Mad2 such that its extended N-terminus drapes into the hexamer pore for engagement and remodeling (Alfieri et al., 2018; Ye et al., 2017). In our structure of *Ec* CdnC:HORMA:Pch2^EQ^, the *Ec* Pch2^EQ^ hexamer adopts a right-handed spiral conformation, with four subunits (monomers A-D) bound to ATP, and two subunit (monomers E and F) unbound (Fig. S8A-B). The four ATP-bound subunits overlay closely, while monomers E and F show rotation of the large and small AAA subdomains with respect to one another (Fig. S8C). Monomer F is positioned at the “bottom” of the spiral, and as such shows the most significant conformational differences compared to the other five subunits. Overall, the structure of the *Ec* Pch2^EQ^ hexamer closely resembles that of other substrate-engaged AAA+ ATPases, including mammalian TRIP13 (Alfieri et al., 2018).

As noted above, eukaryotic TRIP13 recognizes its HORMA domain substrate Mad2 through the adapter protein p31^comet^ (Alfieri et al., 2018; Ye et al., 2017). Our structure of *Ec* CdnC:HORMA:Pch2^EQ^ reveals that in this complex, CdnC serves as the adapter, binding the top face of the Pch2 hexamer at the interface of monomers E and F (Fig. 5A). CdnC-Pch2 binding positions *Ec* HORMA over the Pch2 hexamer pore, with its extended N-terminus draping into the pore and interacting with the pore loops from Pch2 monomers A-E (Fig. 5B). These pore loops form a tight spiral and engage the extended HORMA N-terminus with two-residue periodicity: Pch2 pore loop 1 residues Gly119 and Val121 in monomers A-E engage even-numbered residues from HORMA (Ser2, Tyr4, Tyr6, Val8, and Glu10) through main-chain hydrogen bonding (Fig. 5C-D). At the same time, conserved histidine residues (His173) in pore loop 2 of Pch2 monomers A-D form a “ladder” with each histidine side-chain forming two hydrogen bonds with the side-chains of serine/threonine residues at odd-numbered positions in the HORMA N-terminus (Ser3, Ser5, and Thr7) (Fig. 5C-D). In keeping with these highly specific interactions, we observe a striking pattern of conservation in the N-termini of HORMA proteins from 1-HORMA operons, with conserved serine/threonine residues in every second position (Fig. 5E). This conservation pattern extends to *Ec* HORMA residue 17, strongly suggesting that *Ec* Pch2 can undergo at least 3-4 cycles of ATP hydrolysis and translocation before disengaging, sufficient to partially unfold helix αA and disrupt both HORMA-CdnC binding (which primarily involves helix aA) and HORMA-closure motif binding (by destabilizing the C-terminal safety belt). Thus, our structure of the *Ec* CdnC:HORMA:Pch2^EQ^ complex supports a model in which Pch2 disassembles a CD-NTase:HORMA complex by remodeling the HORMA domain protein, and provides the highest-resolution view to date of a AAA+ ATPase hexamer engaged with its substrate.

Our biochemical data shows that addition of Pch2 inactivates second messenger synthesis by *Ec* CdnC+HORMA+DNA, with Pch2 effectively inhibiting second messenger synthesis at a 1:60 ratio of Pch2 hexamer to CdnC+HORMA (Fig. 1C, sample vi). This data is consistent with the idea that Pch2 catalytically disassembles and inactivates the CdnC:HORMA complex, likely through engagement and unfolding of the HORMA N-terminus. In agreement with this idea, we found that Pch2 is unable to effectively inhibit second messenger synthesis when ATP binding is disrupted by a mutation to the Walker A motif (K87A; Fig. 5F, sample iii). Further, Pch2 is unable to inhibit second messenger synthesis by an *Ec* CdnC:HORMA-ΔN complex, which lacks the disordered N-terminus of *Ec* HORMA and thus cannot be engaged by Pch2 (Fig. 5F, samples iv-v). These data support a model in which bacterial Pch2 enzymes attenuate signaling by their cognate CD-NTases by disassembling an active CD-NTase:HORMA complex.

### NucC is a cAAA-activated DNA endonuclease

Bacterial CD-NTase+HORMA+Pch2 operons encode one of several putative effector proteins, which include proteases, phospholipases, and at least three families of putative endo- and exonucleases (Burroughs et al., 2015). Both the *E. coli* MS115-1 and *P. aeruginosa* ATCC27853 operons encode a restriction endonuclease-related protein we term NucC (<u>Nuc</u>lease, <u>C</u>D-NTase associated). We purified *E. coli* MS115-1 NucC and tested its ability to degrade purified plasmid DNA in the presence of second messengers including cyclic and linear di-AMP, and the cyclic trinucleotides cAAA and cAAG. We found that NucC degrades DNA to ~50-100 bp fragments in the presence of low-nanomolar concentrations of cAAA, and is also activated to a lesser extent by cyclic di-AMP and cAAG (Fig. 6A). NucC is apparently insensitive to DNA methylation: the enzyme is equally active on unmethylated PCR product, plasmid purified from a K-12 based *E. coli* strain (NovaBlue), and plasmid purified from strain MS115-1, which harbors the CdnC operon and several predicted DNA methylases not found in K-12 strains (Fig. S9A). Thus, NucC is DNA endonuclease activated by cAAA, the second messenger product of its cognate CD-NTases.

**Figure 6.**
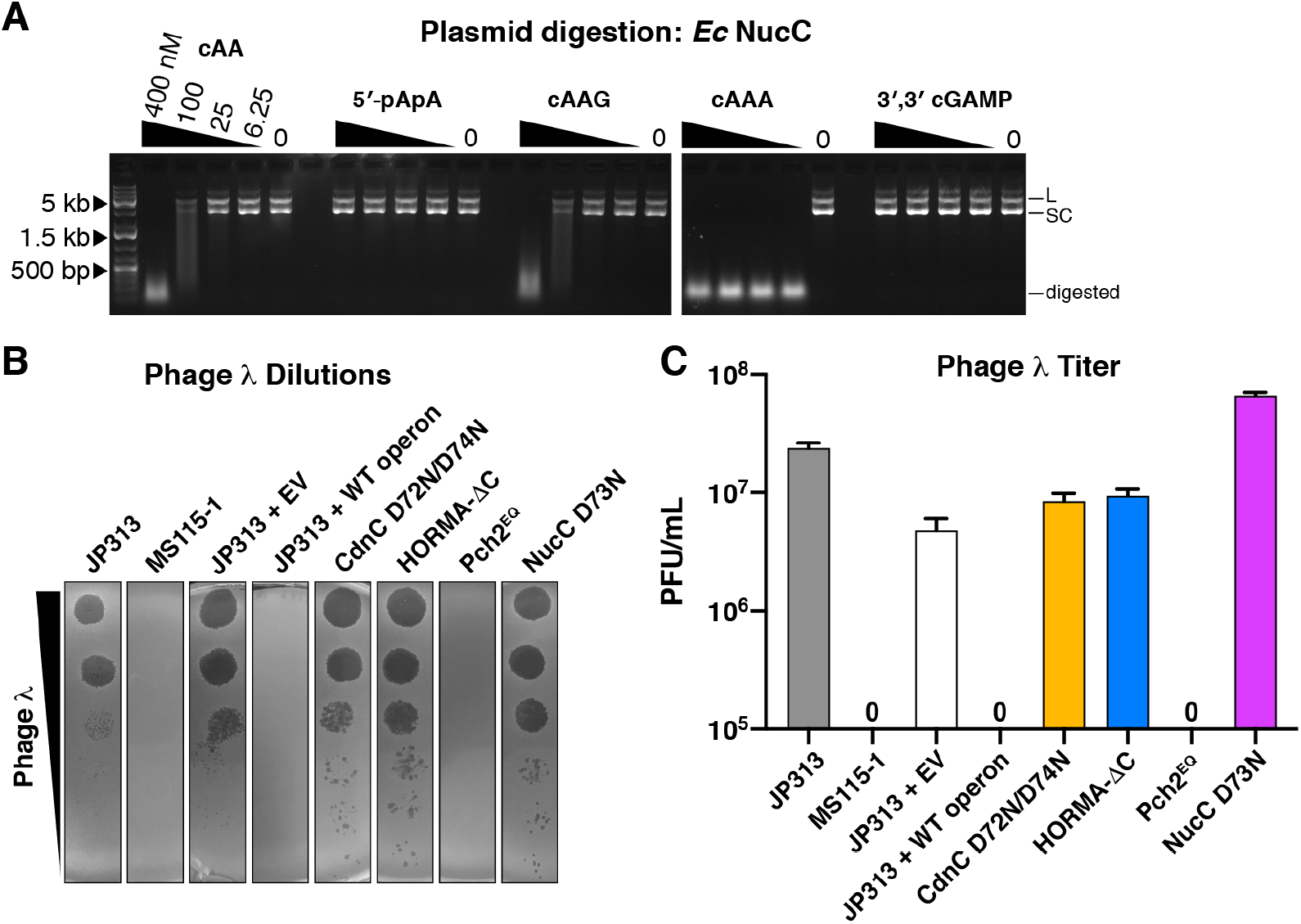
The *E. coli* MS115-1 CD-NTase operon confers bacteriophage immunity through activation of a DNA endonuclease. **(A)** Plasmid digestion assay with Ec NucC (10 nM) and the indicated concentrations of cyclic and linear second messengers. “L” denotes linear plasmid, “SC” denotes closed-circular supercoiled plasmid, and “cut” denotes fully-digested DNA. **(B)** Dilution of bacteriophage λ on lawns of *E. coli* JP313 (wild-type laboratory strain), MS115-1, and JP313+plasmid-encoded *E. coli* MS115-1 CD-NTase operons (blue) with wild-type proteins (WT) or the indicated mutations. EV: empty vector. Six 10-fold bacteriophage dilutions are shown. **(C)** Quantitation of bacteriophage λ infectivity. Shown is the average +/− standard deviation of three trials at a single bacteriophage dilution (representative plate images in Fig. S9B). The three strains marked “0” showed no plaques, indicating infectivity is < 1×10^5^ PFU/mL for these strains.

### CD-NTase operons confer bacteriophage immunity onto their bacterial host

Our structural and biochemical data suggest that bacterial CD-NTases and HORMA domain proteins constitute a “sensor module” that recognizes specific peptides from a foreign invader, assembles an active CD-NTase:HORMA complex to produce a cAAA second messenger, thereby activating a defensive response by the effector nuclease NucC. To directly test the idea that these operons serve as a defense against bacteriophages, we first tested *E. coli* MS115-1, which harbors the CdnC operon, for resistance to bacteriophage λ. In contrast to a laboratory strain of *E. coli* (JP313; (Economou et al., 1995)), *E. coli* MS115-1 is immune to bacteriophage λ infection as judged by plaque formation (Fig. 6B-C). We next inserted the entire *E. coli* MS115-1 operon (including CdnC, HORMA, Pch2, and NucC; see Supplemental Note) into *E. coli* JP313 on a plasmid vector, and found that the operon confers robust immunity to phage λ (Fig. 6B-C). Immunity was compromised when we disrupted second messenger synthesis by CdnC (CdnC D72N/D74N mutant), HORMA-CdnC binding (HORMA-ΔC mutant), or NucC’s DNA cleavage activity (NucC D73N mutant; based on sequence alignments with type II restriction endonucleases), demonstrating that immunity depends on both the sensor and effector modules in this pathway (Fig. 6B-C). Disruption of Pch2’s ATP hydrolysis activity (Pch2^EQ^ mutant) did not compromise immunity, consistent with the idea that Pch2 is a negative regulator of this pathway (Fig. 6B-C). Thus, CD-NTase+HORMA+Pch2+NucC operons constitute a new bacteriophage resistance pathway that is broadly distributed among bacteria, including important human pathogens like *E. coli* and *P. aeruginosa*.

## Discussion

The recent discovery and classification of a diverse family of cGAS/DncV-like nucleotidyltransferases (CD-NTases) that are sparsely distributed among important environmental and pathogenic bacteria raised many important questions about these proteins’ molecular mechanisms and the biological functions of their associated operons (Burroughs et al., 2015; Whiteley et al., 2019). Here, we address these questions and define the functions of a group of CD-NTase operons encoding eukaryotic-like HORMA domain proteins and Pch2-like ATPase regulators. We define the molecular mechanisms of CD-NTase activation and inactivation, identify the second messenger products of these enzymes, and show that second messenger production activates the effector nuclease NucC. Together, these proteins constitute a pathway that confers robust bacteriophage immunity onto its bacterial host.

Our data support a model for HORMA/Pch2 regulation of CD-NTase activity in bacteria that is remarkably reminiscent of the roles of Mad2 and Pch2/TRIP13 in the eukaryotic spindle assembly checkpoint. In this checkpoint pathway, the HORMA domain protein Mad2 is kept in its inactive open state through constant low-level conformational conversion by Pch2/TRIP13 (Kim et al., 2018; Ma and Poon, 2016, 2018). Upon entrance into mitosis, large-scale assembly of Mad2 into a “mitotic checkpoint complex” with the closure motif-bearing protein Cdc20 overwhelms the disassembly capacity of Pch2/TRIP13, resulting in checkpoint activation. After successful kinetochore-microtubule attachment and cessation of mitotic checkpoint complex assembly, the checkpoint is inactivated as existing complexes are disassembled by Pch2/TRIP13 and an alternative pathway (Kim et al., 2018). In a similar manner, we propose that in the absence of a foreign threat, bacterial HORMA domain proteins are maintained in the inactive “open” state by continual activity of Pch2 (Fig. 7, step 1), thereby inhibiting CD-NTase activation and second messenger synthesis. Introduction of specific phage proteins into the cell upon infection or lysogenic induction, followed by HORMA domain protein recognition, binding, and conversion to the closed state, results in CD-NTase activation and cAAA synthesis (Fig. 7, step 2). cAAA in turn activates the effector nuclease NucC, which may specifically destroy the invading phage genome, or alternatively destroy the host genome and cause cell death in an abortive infection mechanism (Fig. 7, step 3). In the former case, Pch2 would then mediate disassembly and inactivation of CD-NTase:HORMA complexes after elimination of the phage threat (Fig. 7, step 4).

**Figure 7.**
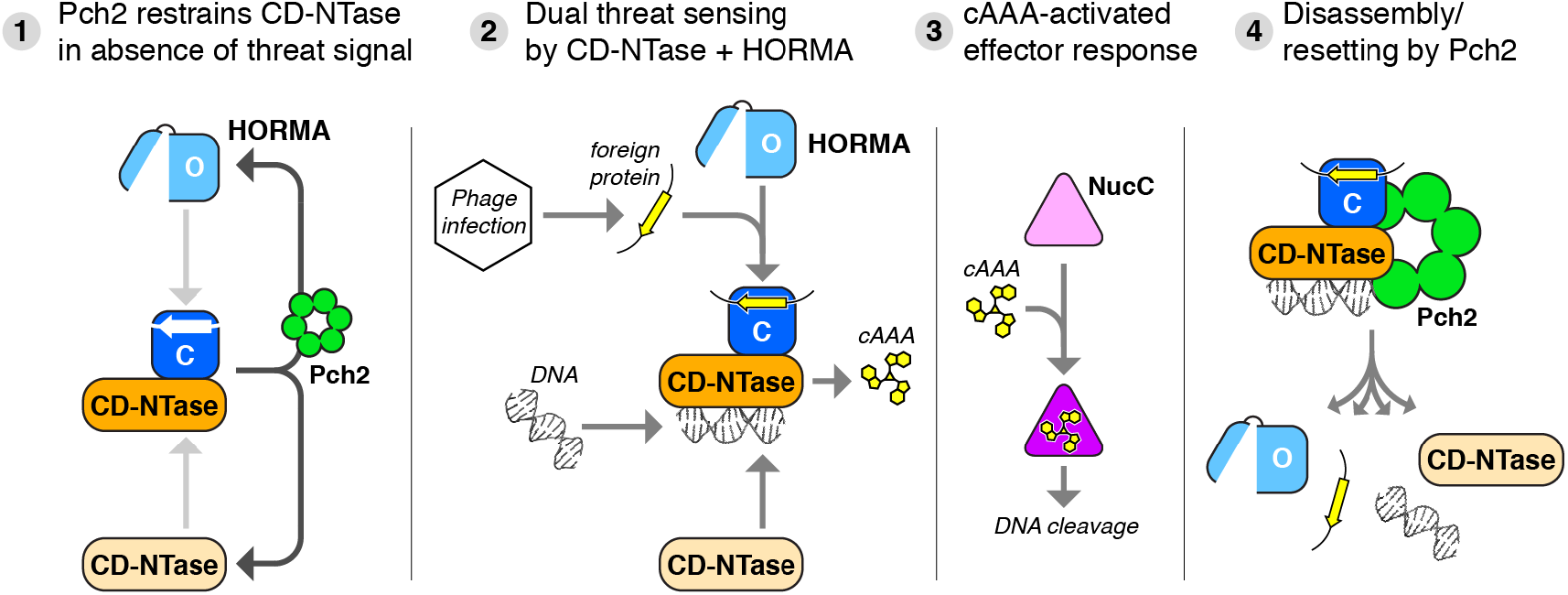
Model for bacteriophage sensing and immunity by CD-NTase+HORMA+Pch2 operons. (1) In the absence of a bacteriophage threat, Pch2 restrains CD-NTase activation by converting closed HORMA proteins (dark blue) to the open state (light blue). (2) Upon infection, open HORMA proteins recognize closure motif sequences in foreign proteins (yellow), convert to the closed state, then bind their cognate CD-NTase (orange). The CD-NTase:HORMA complex (which further requires bound DNA in some cases) synthesizes a nucleotide-based second messenger (cAAA). (3) Second messengers bind and activate effector proteins including NucC (Lau et al., 2019). (4) After elimination of the infection, Pch2 disassembles the active CD-NTase:HORMA complex to attenuate signaling and reset HORMA into the open state.

While we demonstrate the structural mechanisms of CD-NTase activation by the first-identified bacterial HORMA domain proteins, important questions remain regarding these proteins’ evolution and function. First, it will be important to identify the closure motif-containing proteins recognized by bacterial HORMA domain proteins. The consensus closure motif sequences we identified by phage display do not allow clear identification of likely targets by sequence searches, so more direct tests will be required to identify these targets. The operon’s utility to bacterial survival would be maximized if the HORMA proteins were to recognize a widely-conserved bacteriophage protein, rather than a protein that is only found in a few bacteriophages. It would also be beneficial to recognize phage infection as early as possible, potentially by recognizing initial invasion through tail spike detection, or by recognizing a protein that is transcribed and translated early in the phage infection cycle.

Our data also do not address the roles of HORMA1 versus HORMA2 in bacterial 2-HORMA operons. We show that *Pa* HORMA2 binds closure motifs and activates *Pa* CdnD, and is therefore functionally equivalent to *Ec* HORMA. *Pa* HORMA1, meanwhile, does not bind closure motifs and appears to be locked in the open conformation. We show that HORMA1 is able to bind both CdnD and HORMA2, and thereby scaffold a larger CdnD:HORMA1:HORMA2 complex, but the exact role of this larger complex remains unknown.

Another key question is how eukaryotic-like HORMA and Pch2 proteins have evolved to function in bacteria, in concert with a cGAS-like nucleotidyltransferase. Evolutionary studies have shown that the last eukaryotic common ancestor (LECA) possessed at least one HORMA domain protein and an ancestor of TRIP13 (van Hooff et al., 2017), but neither HORMA domain proteins nor TRIP13 have previously been reported in bacteria. We propose that these proteins were obtained by an ancestral bacterium or virus by horizontal gene transfer from a eukaryotic host, and were subsequently co-opted to function in second messenger signaling. Our evolutionary analysis of Pch2/TRIP13 proteins indicate that Pch2 homologs from 2-HORMA CD-NTase operons are more closely related to eukaryotic TRIP13 than their relatives in 1-HORMA operons (Fig. S8G), suggesting that 1-HORMA operons may have evolved from 2-HORMA operons.

Well-characterized CD-NTases including bacterial DncV and mammalian cGAS catalyze cyclic dinucleotide synthesis by binding two nucleotides, and catalyzing nucleophilic attack of each nucleotide’s 2′ or 3′ ribose oxygen by the other nucleotide’s α-phosphate. OAS proteins, on the other hand, synthesize linear 2′-5′ linked oligoadenylates through a similar mechanism that likely involves a single nucleophilic attack, followed by translocation and binding/attack by another ATP (Lohöfener et al., 2015). In this context, a key unanswered question is how bacterial CD-NTases in clades C and D – including *Ec* CdnC, *Pa* CdnD, and *E. cloacae* CdnD02 (Whiteley et al., 2019) – catalyze specific synthesis of cyclic trinucleotide second messengers. The enzymes’ active sites likely cannot accommodate three nucleotides, necessitating translocation of a linear intermediate during synthesis prior to the final cyclization reaction. Both *Ec* CdnC and *Pa* CdnD synthesize cAA as a minor product, suggesting that the enzymes can cyclize both 2-base or 3-base linear intermediates. Additional studies will be required to understand the mechanism and specificity of this reaction.

Here, we have defined the molecular mechanisms and biological role of a class of bacterial CD-NTase operons with associated HORMA and Pch2-like regulators and the nuclease effector NucC. Bacterial CD-NTase operons are extraordinarily diverse, however, with multiple regulatory/sensor systems including the HORMA/Pch2 systems described here and a putative ubiquitin-conjugation system, and a variety of putative effectors including nucleases, proteases, and phospholipases (Burroughs et al., 2015). Thus, the spectrum of biological functions for bacterial CD-NTases, and the potential for cross-kingdom signaling through recognition of bacterial second messengers by eukaryotic host proteins, is only just beginning to be explored.

## Materials and Methods

### Yeast Two-hybrid

Coding sequences for genes from operon 300414846 from *E. coli* MS115-1 (CD-NTase018 from (Whiteley et al., 2019)) were synthesized (GeneArt) and inserted into vectors pBridge and pGADT7 (Clontech) by isothermal assembly. Coding sequence for genes from *P. aeruginosa* strain 27853 (CD-NTase023 from (Whiteley et al., 2019)) were amplified from genomic DNA (American Type Culture Collection) and inserted into vectors pBridge and pGADT7 by isothermal assembly. See Table S2 for all protein sequences. pBridge vectors were transformed into *S. cerevisiae* strain AH109 and selected on SC media lacking tryptophan (-TRP). pGADT7 AD vectors were transformed into *S. cerevisiae* strain Y187 and selected on SC media lacking leucine (-LEU). Haploid strains were mated and diploids selected on SC-TRP/-LEU. Diploid cells were diluted in water and replated onto SC-TRP/-LEU (control), -TRP/-LEU/-HIS (histidine) (low stringency), and -TRP/-LEU/-HIS/-ADE (adenine) (high stringency), grown for 2-3 days, then examined for growth.

For yeast three-hybrid assays, pBridge vectors containing one protein in MCS I were further modified by NotI cleavage at the MCS II site followed by isothermal assembly-mediated insertion of a second gene, resulting in a single vector encoding two genes, with the Gal4-BD tag fused to the N-terminus of the gene in MCS I. These vectors were transformed into AH109 and mated with pGADT7 AD vectors encoding other proteins.

### Protein Expression, Purification, and Characterization

All proteins were cloned into UC Berkeley Macrolab vectors 2AT (for untagged expression) or 2BT (encoding an N-terminal TEV protease-cleavable His_6_-tag). Co-expression cassettes were assembled by amplifying genes from these vectors by PCR, and re-inserting into vector 2BT so that one protein is tagged. For coexpression of *Pa* HORMA2 with Peptide 1, the peptide sequence (EVMEFNP) was cloned into UC Berkeley Macrolab vector 2CT (encoding an N-terminal TEV protease-cleavable His_6_-maltose binding protein tag), then assembled into coexpression vectors with *Pa* HORMA2, or *Pa* HORMA2 and CdnD, or *Pa* HORMA2 and HORMA1.

Proteins were expressed in *E. coli* strain Rosetta 2 (DE3) pLysS (EMD Millipore, Billerica MA). Cultures were grown at 37°C to A600=0.5, then induced with 0.25 mM IPTG and shifted to 20°C for 15 hours. Cells were harvested by centrifugation and resuspended in buffer A (20 mM Tris pH 7.5 (Tris pH 8.5 for CdnC/CdnD), 10% glycerol) plus 400 mM NaCl, 10 mM imidazole, and 5 mM β-mercaptoethanol. Proteins were purified by Ni^2+^-affinity (Ni-NTA agarose, Qiagen) then passed over an anion-exchange column (Hitrap Q HP, GE Life Sciences, Piscataway NJ) in Buffer A plus 100 mM NaCl and 5 mM β-mercaptoethanol, collecting flow-through fractions. Tags were cleaved with TEV protease (Tropea 2009), and cleaved protein was passed over another Ni^2+^ column (collecting flow-through fractions) to remove uncleaved protein, cleaved tags, and tagged TEV protease. The protein was passed over a size exclusion column (Superdex 200, GE Life Sciences) in buffer GF (buffer A plus 300 mM NaCl and 1 mM dithiothreitol (DTT)), then concentrated by ultrafiltration (Amicon Ultra, EMD Millipore) to 10 mg/ml and stored at 4°C. For selenomethionine derivatization, protein expression was carried out in M9 minimal media supplemented with amino acids plus selenomethionine prior to IPTG induction (Van Duyne et al., 1993), and proteins were exchanged into buffer containing 1 mM tris(2-carboxyethyl)phosphine (TCEP) after purification to maintain the selenomethionine residues in the reduced state.

For characterization of oligomeric state by size exclusion chromatography coupled to multi-angle light scattering (SEC-MALS), 100 μL of purified protein/complex at 2-5 mg/mL was injected onto a Superdex 200 Increase 10/300 GL or Superose 6 Increase 10/300 GL column (GE Life Sciences) in a buffer containing 20 mM HEPES pH 7.5, 300 mM NaCl, 5% glycerol, and 1 mM DTT. Light scattering and refractive index profiles were collected by miniDAWN TREOS and Optilab T-rEX detectors (Wyatt Technology), respectively, and molecular weight was calculated using ASTRA v. 6 software (Wyatt Technology).

### Crystallization and structure determination

#### E. coli CdnC

We obtained crystals of *Ec* CdnC using two different methods. First, small-scale dialysis of CdnC at 10 mg/mL from buffer GF to a buffer containing 0% glycerol and 200 mM NaCl resulted in the formation of large prism crystals. Second, mixing CdnC (10 mg/mL in buffer GF) 1:1 with a well solution containing 20 mM HEPES pH 8.1, 10-12% PEG 4000, 100 mM NaCl, 10 mM MgCl_2_ (plus 1 mM TCEP for selenomethionine-derivatized protein) in hanging-drop format resulted in smaller prism crystals. All crystals were cryoprotected by the addition of 2-Methyl-2,4-pentanediol (MPD; 30% for dialysis crystals, 15% for hanging-drop), and flash-frozen in liquid nitrogen. We collected diffraction data at the Advanced Photon Source NE-CAT beamline 24ID-E (support statement below), and all datasets were processed with the RAPD data-processing pipeline, which uses XDS (Kabsch, 2010) for data indexing and reduction, AIMLESS (Evans and Murshudov, 2013) for scaling, and TRUNCATE (Winn et al., 2011) for conversion to structure factors. We determined the structure by single-wavelength anomalous diffraction methods using a 1.92 Å dataset from selenomethionine-derivatized protein, in the PHENIX Autosol wizard (Terwilliger et al., 2009). We manually rebuilt the initial model in COOT (Emsley et al., 2010), and refined against a 1.44 Å native dataset in phenix.refine (Afonine et al., 2012) using positional and individual anisotropic B-factor refinement (Table S1). Since ATP was not added to Ec CdnC purification or crystallization buffers, we conclude that ATP observed in the *Ec* CdnC active site was co-purified with the enzyme.

#### P. aeruginosa CdnD

Crystals of wild-type *Pa* CdnD (Apo state) were obtained by mixing protein (20 mg/mL) 1:1 with well solution containing 0.1 M HEPES pH 7.5 (or imidazole pH 8.0) and 1.8-2.0 M Ammonium sulfate in hanging-drop format. Crystals were cryoprotected by the addition of 30% glycerol, and flash-frozen in liquid nitrogen. Diffraction data were collected at the Advanced Photon Source beamline 24ID-E and processed with the RAPD data-processing pipeline. Crystals of *Pa* CdnD D62N/D64N (ATP-bound) were obtained by mixing protein (20 mg/mL) in crystallization buffer plus 10 mM MgCl_2_ and 5 mM ATP 1: 1 with well solution containing 0.1 M Tris pH 8.5, 0.2 M sodium formate, and 20-22% PEG 6000 in hanging drop format. Crystals were cryoprotected by the addition of 10% glycerol and flash-frozen in liquid nitrogen. Diffraction data were collected at the Advanced Photon Source beamline 24ID-E and processed with the RAPD data-processing pipeline.

We determined the structure of Apo *Pa* CdnD using a 2.05 Å-resolution single-wavelength dataset from a crystal of selenomethionine-derivatized protein, using the PHENIX Autosol wizard. We manually rebuilt the initial model in COOT, followed by refinement in phenix.refine using positional and individual B-factor refinement. The structure of ATP-bound *Pa* CdnD D62N/D64N was determined by molecular replacement in PHASER (McCoy et al., 2007).

#### P. aeruginosa HORMA1

Crystals of *Pa* HORMA1 were obtained by mixing protein (10 mg/mL) in crystallization buffer 1:1 with well solution containing 100 mM imidazole pH 8.0, 200 mM CaCl_2_, and 32% PEG 3350 in hanging-drop format. Crystals were flash-frozen in liquid nitrogen directly from the crystallization drop. We collected diffraction data at the Advanced Photon Source NE-CAT beamline 24ID-C, and all datasets were processed with the RAPD pipeline. In order to obtain phases by selenomethionine derivatization, we designed several mutants with conserved hydrophobic residues mutated to methionine. One mutant, V102M/L146M, crystallized more robustly than wild-type protein and was used for all later crystallographic analysis. We determined the structure of *Pa* HORMA1 V102M/L146M from a 1.86 Å resolution dataset from crystals derivatized with NaBr (1 minute soak in cryoprotectant solution containing 1 M NaBr). Data was processed with the RAPD pipeline, Br sites were identified with hkl2map (Pape et al., 2004; Sheldrick, 2010), and the structure was determined using the phenix AUTOSOL wizard. The initial models was manually rebuilt with COOT and refined against a 1.64 Å resolution native dataset using phenix.refine.

#### P. aeruginosa HORMA2

We obtained crystals of *Pa* HORMA2 by mixing protein at 10 mg/mL 1:1 in hanging drop format with well solution containing 100 mM Imidazole pH 8.0, 0.8 M NaH_2_PO_4_, and 0.55-0.8 M KH_2_PO_4_. Crystals were cryoprotected with an additional 30% glycerol and flash-frozen in liquid nitrogen. Native data (to 2.1 Å resolution) were collected at APS beamline 24ID-C, and a single-wavelength anomalous diffraction dataset (to 2.5 Å resolution) on a crystal grown from selenomethionine-derivatized protein was collected at SSRL beamline 9-2. Data were processed by the RAPD pipeline (native) or the autoxds pipeline (selenomethionine). The structure was determined by SAD phasing using the PHENIX AutoSol wizard, models were manually rebuilt with COOT, and refined against the native dataset using phenix.refine.

#### P. aeruginosa HORMA2-ΔC

We purified *Pa* HORMA2-ΔC (residues 2-133) as above (replacing Tris-HCl with HEPES pH 8.0 in all buffers), then dimethylated surface lysine residues by mixing protein at 1 mg/mL with 50 mM freshly-prepared dimethylamine borane complex and 100 mM formaldehyde, incubating at 4°C for one hour, then quenching by addition of 25 mM glycine. Dimethylated *Pa* HORMA2-ΔC was then re-concentrated to 17 mg/mL and buffer-exchanged into GF buffer to remove residual reaction components.

We obtained crystals of dimethylated *Pa* HORMA2-ΔC by mixing mixed purified protein at 17 mg/mL 1:1 in hanging drop format with well solution containing 100 mM Tris-HCl pH 8.5, 10 mM NiCl_2_, and 20% PEG 2000 MME. Crystals were cryoprotected with an additional 10% glycerol or 10% PEG 400, then looped and frozen in liquid nitrogen. Diffraction data were collected at APS beamline 24ID-C and processed by the RAPD pipeline. We determined the structure by molecular replacement using the structure of *Pa* HORMA2 as a search model (the successful model lacked the C-terminal safety belt region and the N-terminus, which both undergo conformational changes between the open and closed states. Initial models were manually rebuilt with COOT and refined using phenix.refine.

#### P. aeruginosa HORMA1:HORMA2:Peptide 1

A complex of *Pa* HORMA1 (residues 1-133), HORMA2, and Peptide 1 (EVMEFNP) was obtained by co-expressing and purifying His_6_-MBP-tagged Peptide 1 with untagged HORMA1 and HORMA2, with the tag cleaved prior to crystallization. Crystals were obtained by mixing protein (10 mg/mL) in hanging-drop format 1:1 with well solution containing 100 mM HEPES pH 7.5, 0.5 M sodium acetate, 31% PEG 3350, and 1 mM ATP. Crystals were flash-frozen in liquid nitrogen directly from the crystallization drop. We collected a 2.0 Å diffraction dataset on SSRL beamline 9-2, processed the data using the autoxds pipeline, and determined the structure by molecular replacement with PHASER, using the structures of *Pa* HORMA1 and *Pa* HORMA2. Initial models were manually rebuilt with COOT and refined using phenix.refine.

#### Rhizobiales sp. Pch2

*Rh* Pch2 was purified as above, except all purification buffers contained 5 mM MgCl_2_. We obtained crystals of selenomethionine-derivatized *Rh* Pch2 by mixing protein at 24 mg/mL 1:1 in hanging drop format with well solution containing 100 mM sodium citrate pH 6.0, 1.6 M NH_2_SO_4_, and 0.2 M sodium/potassium tartrate. Crystals were cryoprotected with 30% glycerol, and flash-frozen in liquid nitrogen. We collected a 2.05 Å-resolution single-wavelength anomalous diffraction dataset on APS beamline 24ID-E, processed the data with the RAPD pipeline, and determined the structure using the PHENIX Autosol wizard. Initial models were manually rebuilt with COOT and refined using phenix.refine. The structure of *Rh* Pch2 is shown in Fig. S7.

#### P. aeruginosa HORMA2:Peptide 1:CdnD

We obtained crystals of *Pa* HORMA2:Peptide 1:CdnD by mixing purified complex at 15 mg/mL 1:1 in hanging drop format with well solution containing 100 mM Bis-Tris pH 5.5, 0.2 M ammonium acetate, and 30% PEG 3350. Crystals were flash-frozen in liquid nitrogen directly from the crystallization drop. Diffraction data were collected at APS beamline 24ID-C, and processed by the RAPD pipeline. The structure was determined by molecular replacement in PHASER using the structures of *Pa* HORMA2 and CdnD as search models. Initial models were manually rebuilt with COOT, and refined using phenix.refine.

#### E. coli HORMA:CdnC:Pch2^EQ^ complex

To assemble a HORMA:CdnC:Pch2^EQ^ complex, we separately purified *Ec* HORMA:CdnC (coexpressed, with an N-terminal His_6_-tag on HORMA) and *Ec* Pch2^EQ^ (E159Q), then mixed at a ratio of one 1:1 HORMA:CdnC complex per Pch2^EQ^ hexamer in buffer GF plus 1 mM ATP and 5 mM MgCl_2_ at 4°C. After a one-hour incubation, the mixture was passed over a Superdex 200 size exclusion column (GE Life Sciences), and assembled complexes were collected and concentrated by ultrafiltration. Crystals were obtained by mixing protein at 10 mg/mL 1:1 with well solution containing 0.1 M MOPS pH 7.0, 0.2 M MgCl_2_, and 14% PEG 3350 in hanging-drop format. We collected a 2.64 Å resolution dataset at APS beamline 24ID-E, and processed the data using the RAPD pipeline. Crystals adopt space group P2_1_, with one 1 HORMA, 1 CdnC, and 6 Pch2^EQ^ monomers per asymmetric unit. We determined the structure by molecular replacement in PHASER, using the structure of *Ec* CdnC and a Pch2 hexamer model built by separately overlaying the large and small AAA subdomains of six *Rh* Pch2 monomers onto a recent cryo-EM structure of human TRIP13 bound to p31^comet^:MAD2 (Alfieri et al., 2018). A model of *Pa* HORMA2 was used as a guide to manually build *Ec* HORMA into difference density obtained from initial refinement. Initial models were manually rebuilt with COOT and refined using phenix.refine.

### APS NE-CAT Support Statement

This work is based upon research conducted at the Northeastern Collaborative Access Team beamlines, which are funded by the National Institute of General Medical Sciences from the National Institutes of Health (P41 GM103403). The Eiger 16M detector on 24-ID-E beam line is funded by a NIH-ORIP HEI grant (S10OD021527). This research used resources of the Advanced Photon Source, a U.S. Department of Energy (DOE) Office of Science User Facility operated for the DOE Office of Science by Argonne National Laboratory under Contract No. DE-AC02-06CH11357.

### SSRL Support Statement

Use of the Stanford Synchrotron Radiation Lightsource, SLAC National Accelerator Laboratory, is supported by the U.S. Department of Energy, Office of Science, Office of Basic Energy Sciences under Contract No. DE-AC02-76SF00515. The SSRL Structural Molecular Biology Program is supported by the DOE Office of Biological and Environmental Research, and by the National Institutes of Health, National Institute of General Medical Sciences (including P41GM103393). The contents of this publication are solely the responsibility of the authors and do not necessarily represent the official views of NIGMS or NIH.

### Isothermal Titration Calorimetry

Isothermal titration calorimetry was performed on a MicroCal iTC200 in buffer GF, by injecting ATP or GTP (1 mM) into a solution containing 0.1 mM Pa CdnD (wild-type or D62N/D64N mutant). Two injections were performed per protein/nucleotide combination.

### Phage Display

For phage display, we used a commercial assay kit (Ph.D.-7 Phage Display Peptide Library Kit, New England Biolabs) and followed the recommended protocol for “solution phase panning with affinity bead capture” with the following modifications. We used His_6_-tagged *Ec* HORMA, *Pa* HORMA1, and *Pa* HORMA2 as bait proteins. For affinity purification, we diluted 2 μL of bait protein at 0.2 mM and 10 μL phage library into 200 μL volume in TBST (50mM Tris pH7.5, 150mM NaCl, plus 0.1% Tween (first round) or 0.5% Tween (subsequent rounds)). After a 20 minute incubation at room temperature, 50 μL of magnetic Ni-NTA beads (Qiagen; pre-blocked with blocking buffer (0.1M NaHCO_3_, pH 8.6, 5mg/mL BSA, 0.02% NaN_3_) and washed 3x with TBST) were added, and the mixture was incubated a further 15 minutes at room temperature. Beads were washed 10x with wash buffer (50 mM Tris-HCl pH 7.5, 300 mM NaCl, plus 0.1% Tween (first round) or 0.5% Tween (subsequent rounds)). For elution, 1 mL elution buffer (0.2M Glycine-HCl, pH 2.2, 1mg/mL BSA) was added, incubated for 10 minutes at room temperature, then neutralized the eluate by adding 150 μL Tris-HCl pH 9.1. Eluted phage were titered and amplified according to standard protocols, and the selection was repeated, Following four rounds of selection, phages were isolated and the variable peptide region sequenced for at least 20 individual clones.

### Second messenger synthesis

Cyclic trinucleotide synthesis was performed essentially as described (Whiteley et al., 2019). Briefly, 2 mL synthesis reactions contained 1 μM (for *Ec* CdnC) or 3 μM (for *Pa* CdnD) CD-NTase or CD-NTase:HORMA complex and 0.25 mM ATP (plus 0.25 mM GTP for *Pa* CdnD) in reaction buffer with 12.5 mM NaCl, 20 mM MgCl_2_, 1 mM DTT, and 10 mM Tris-HCl pH 8.5 (or HEPES-NaOH pH 7.5 for reactions with *Pa* Pch2). For reactions with DNA, plasmid DNA (pUC18) was used at 0.15 μg/mL, 4 kb linear DNA was used at 0.1 μg/mL, single-stranded DNA (sheared salmon sperm DNA; Invitrogen #AM9680) was used at 1 μg/mL, and 40 bp dsDNA was used at 13 μg/mL. For reactions with Pch2, *Ec* Pch2 was used at 100 nM monomer/16.67 nM hexamer and *Pa* Pch2 was used at 300 nM monomer/50 nM hexamer; this represents a 60-fold lower molar concentration of active Pch2 hexamers compared to CdnC/HORMA in the reactions. Reactions were incubated at 37°C for 16 hours, then 2.5 units/mL reaction volume (5 units for 2 mL reaction) calf intestinal phosphatase was added and the reaction incubated a further 2 hours at 37°C. The reaction was heated to 65°C for 30 minutes, and centrifuged 10 minutes at 15,000 RPM to remove precipitated protein. Reaction products were separated by ion-exchange chromatography (1 mL Hitrap Q HP, GE Life Sciences) using a gradient from 0.2 to 2 M ammonium acetate. Product peaks were pooled, evaporated using a speedvac, and analyzed by LC-MS/MS.

### Mass spectrometry

To characterize the products of CD-NTase reactions, we performed liquid chromatography-tandem mass spectrometry (LC-MS/MS). LC-MS/MS analysis was performed using a Thermo Scientific Vanquish UHPLC coupled to a Thermo Scientific Q Exactive™ HF Hybrid Quadrupole-Orbitrap™ Mass Spectrometer, utilizing a ZIC-pHILIC polymeric column (100 mm x 2.1 mm, 5 μm) (EMD Millipore) maintained at 45°C and flowing at 0.4 mL/min. Separation of cyclic trinucleotide isolates was achieved by injecting 2 μL of prepared sample onto the column and eluting using the following linear gradient: (A) 20 mM ammonium bicarbonate in water, pH 9.6, and (B) acetonitrile; 90% B for 0.25 minutes, followed by a linear gradient to 55% B at 4 minutes, sustained until 6 minutes. The column was re-equilibrated for 2.50 minutes at 90% B.

Detection was performed in positive ionization mode using a heated electrospray ionization source (HESI) under the following parameters: spray voltage of 3.5 kV; sheath and auxiliary gas flow rate of 40 and 20 arbitrary units, respectively; sweep gas flow rate of 2 arbitrary units; capillary temperature of 275°C; auxiliary gas heater temperature of 350 °C. Profile MS1 spectra were acquired with the following settings; mass resolution of 35,000, AGC volume of 1×10^6^, maximum IT of 75 ms, with a scan range of 475 to 1050 m/z to include z=1 and z=2 ions of cyclic trinucleotides. Data dependent MS/MS spectra acquisition was performed using collision-induced dissociation (CID) with the following settings: mass resolution of 17,500; AGC volume of 1×10^5^; maximum IT of 50 ms; a loop count of 5; isolation window of 1.0 m/z; normalized collision energy of 25 eV; dynamic exclusion was not used. Data reported is for the z=1 acquisition for each indicated cyclic trinucleotide.

### Nuclease Assays

For all nuclease assays, UC Berkeley Macrolab plasmid 2AT (Addgene #29665; 4,731 bp) was used. *Ec* NucC (10 nM unless otherwise indicated) and second messenger molecules were mixed with 1 μg plasmid DNA or PCR product in a buffer containing 10 mM Tris-HCl (pH 7.5), 25 mM NaCl, 10 mM MgCl_2_, and 1 mM DTT (50 μL reaction volume), incubated 10 min at 37°C, then separated on a 1.2% agarose gel. Gels were stained with ethidium bromide and imaged by UV illumination. Second messenger molecules were either purchased from Invivogen or synthesized enzymatically by *Enterobacter cloacae* CdnD02 and purified as previously described (Lau et al., 2019; Whiteley et al., 2019).

### Plaque Assays

To measure bacteriophage immunity, the intact CD-NTase operon from *E. coli* MS115-1 (See Supplemental Note) was PCR-amplified and inserted into vector pLAC22 for IPTG-inducible expression, then modified by PCR-based mutagenesis to generate mutant/tagged protein variants. Vectors were transformed into *E. coli* strain JP313 (Economou et al., 1995) for plaque assays.

For bacteriophage infection plaque assays, a single bacterial colony was picked from a fresh LB agar plate and grown in LB broth containing proper selection antibiotic at 37°C to an OD_600_ of 0.1-0.2. CdnC operon expression was induced through the addition of 0.2 mM IPTG, followed by further growth for one hour to an OD_600_ of 0.6-0.7. For phage dilution assays, 500 μL of cells were mixed with 4.5 mL of of 0.35% LB top agar was added and entire sample was poured on LB plates containing appropriate selection and/or IPTG (0.1 mM). Plates were spotted with 5 μL of bacteriophage λ*cI* (cI gene deleted to inhibit lysogeny and promote clear plaque formation) diluted in phage buffer (10 mM Tris, pH 7.5, 10 mM MgCl_2_, 68 mM NaCl) plus 1 mM CaCl_2_ at six dilutions: ~2.4×10^6^ PFU/mL and five 10-fold dilutions thereof. Plates were incubated at 37°C for 18 hours, then imaged. For plaque numbers, 500 μL of cells at OD_600_ 0.6-0.7 were mixed with 10 μL of bacteriophage λ*cI* (stock concentration ~2.4×10^7^ PFU/mL as measured by infection of JP313, diluted 1000-fold prior to mixing) diluted in phage buffer (10 mM Tris, pH 7.5, 10 mM MgCl_2_, 68 mM NaCl) plus 1 mM CaCl_2_, and incubated at room temperature for 30 min. 4.5 mL of 0.35% LB top agar was added and entire sample was poured on LB plates containing appropriate selection and IPTG (0.1 mM) and incubated at 37°C. Plaques were counted after 18 hours of growth.

## End Matter

## Author Contributions and Notes

Conceptualization, QY and KDC; Methodology, QY, ITM, JDW, MJ, KDC; Investigation, QY, RKL, ITM, JDW, CSA, KDC; Writing – Original Draft, QY, RKL, KDC; Writing – Review & Editing, QY, RKL, MJ, KDC; Visualization, QY, ITM, JDW, KDC; Supervision, MJ and KDC; Project Administration, KDC; Funding Acquisition, MJ and KDC.

The authors declare no conflict of interest.

## Acknowledgments

The authors thank the staffs of the Stanford Synchrotron Radiation Lightsource and the Advanced Photon Source NE-CAT beamlines for assistance with crystallographic data collection; A. Bobkov (Sanford Burnham Prebys Medical Discovery Institute, Protein Analysis Core) for assistance with isothermal titration calorimetry; and A. Desai, M. Daugherty, J. Pogliano, and P. Kranzusch for critical reading and helpful suggestions. RKL was supported by the UC San Diego Quantitative and Integrative Biology training grant (NIH T32 GM127235). ITM was supported by NIH T32 HL007444, T32 GM007752 and F31 CA236405. JDW was supported by NIH K01 DK116917 and P30 DK063491. MJ was supported by NIH S10 OD020025 and R01 ES027595. KDC was supported from the Ludwig Institute for Cancer Research and the University of California, San Diego.

---

**Figure S1.**
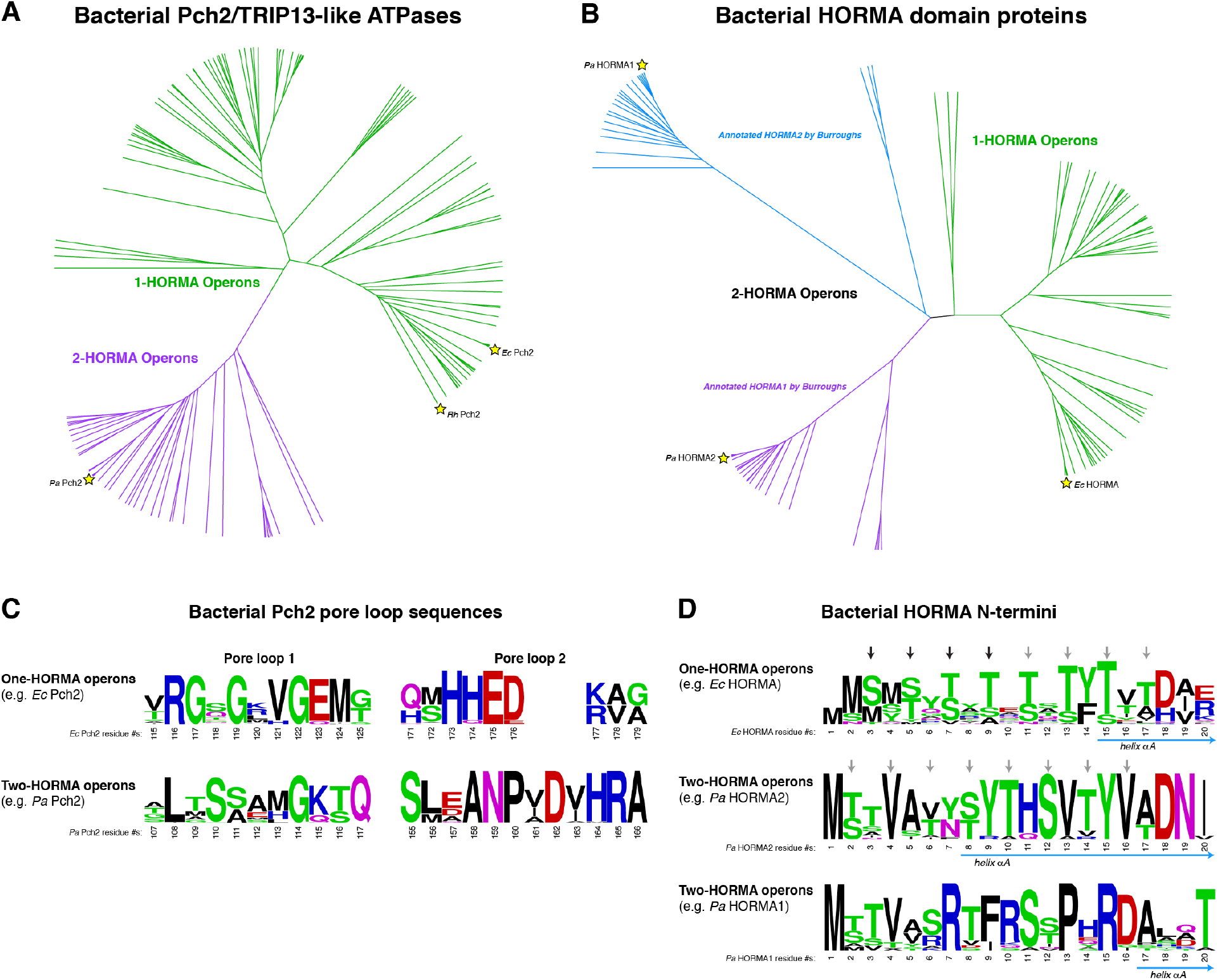
Sequence analysis of bacterial HORMA proteins and Pch2/TRIP13-like ATPases. **(A)** Phylogenetic tree of bacterial Pch2/TRIP13-like ATPases, with proteins from one-HORMA operons colored green, and proteins from two-HORMA operons colored purple. Yellow stars indicate proteins used in this study: *Ec* Pch2, *Pa* Pch2, and *Rhizobiales* (*Rh*) Pch2. **(B)** Phylogenetic tree of bacterial putative HORMA domain proteins, with proteins from one-HORMA operons colored green, and proteins from two-HORMA operons colored blue/purple. Stars indicate proteins used in this study: *Ec* HORMA, *Pa* HORMA1, and *Pa* HORMA2. **(C)** Sequence logos showing divergence of pore loop sequences in Pch2 proteins from one-HORMA and two-HORMA operons. **(D)** Sequence logos showing the N-termini of HORMA proteins from one-HORMA operons (top), and from the two families of HORMA proteins from two-HORMA operons (middle/bottom). The first residue of helix αA (based on crystal structures of *Ec* HORMA, *Pa* HORMA1, and *Pa* HORMA2) are indicated by blue arrows. Arrows above the sequence of *Ec* HORMA (top) indicate serine/threonine residues directly engaged by *Ec* Pch2 pore loop 2 residue His173: black arrows indicate residues engaged by pore loops in our structure, and gray arrows indicate continuation of the conserved pattern into helix αA.

**Figure S2.**
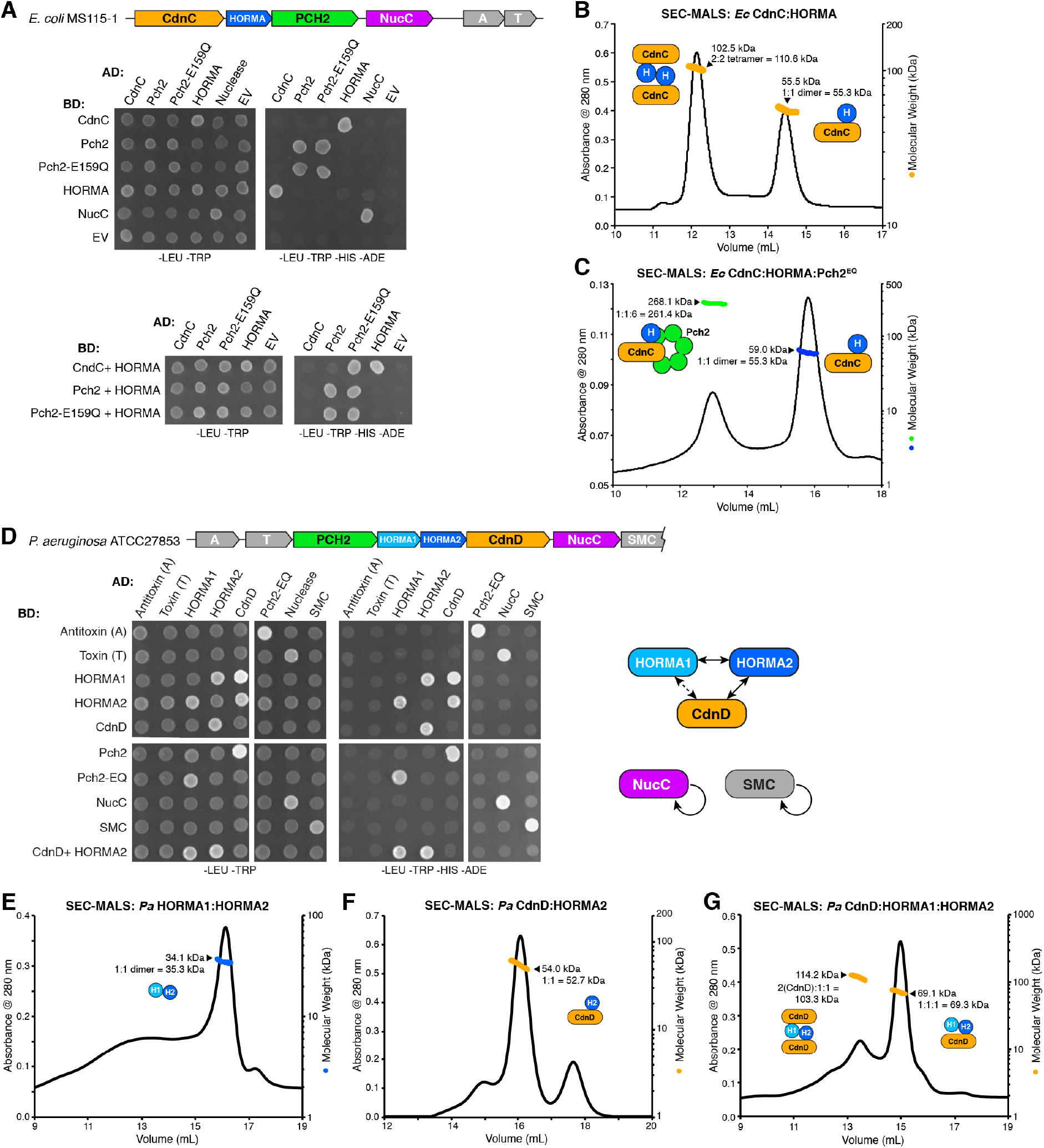
Protein-protein interactions in HORMA/Pch2-associated CD-NTase operons. **(A)** Yeast two-hybrid and three-hybrid analysis of protein-protein interactions in the *E. coli* MS115-1 CD-NTase containing operon. “-LEU-TRP” refers to non-selective growth media, and “-LEU-TRP-HIS-ADE” refers to selective growth media. This operon encodes a RES-Xre type toxin-antitoxin gene pair (Makarova et al. 2009; Skjerning et al. 2018), labeled A/T, which was not tested for protein-protein interactions. “EV” refers to empty vector controls. **(B)** SEC-MALS of the *Ec* CdnC:HORMA complex, showing that it forms 1:1 and 2:2 complexes in solution. **(C)** SEC-MALS of *Ec* CdnC:HORMA:Pch2^EQ^, showing that it forms a 1:1:6 complex in solution when pre-incubated with ATP. **(D)** Yeast two-hybrid and three-hybrid analysis of protein-protein interactions in the *P. aeruginosa* ATCC27853 CD-NTase containing operon. In addition to CdnD, two HORMA domain proteins (HORMA1 and HORMA2), Pch2, and NucC, the operon also encodes a putative toxin-antitoxin gene pair (PIN domain protein (T) and helix-turn-helix protein (A)) and a Rad50-like SMC-family protein (SMC). *Right:* Summary of protein-protein interactions in the *P. aeruginosa* ATCC27853 operon. **(E)** SEC-MALS analysis of *Pa* HORMA1:HORMA2, showing that the proteins form a 1:1 complex in solution. **(F)** SEC-MALS analysis of *Pa* CdnD:HORMA2, showing that the proteins form a 1:1 complex in solution. **(G)** SEC-MALS analysis of *Pa* CdnD:HORMA1:HORMA2, showing that the proteins form a 2:1:1 complex in solution.

**Figure S3.**
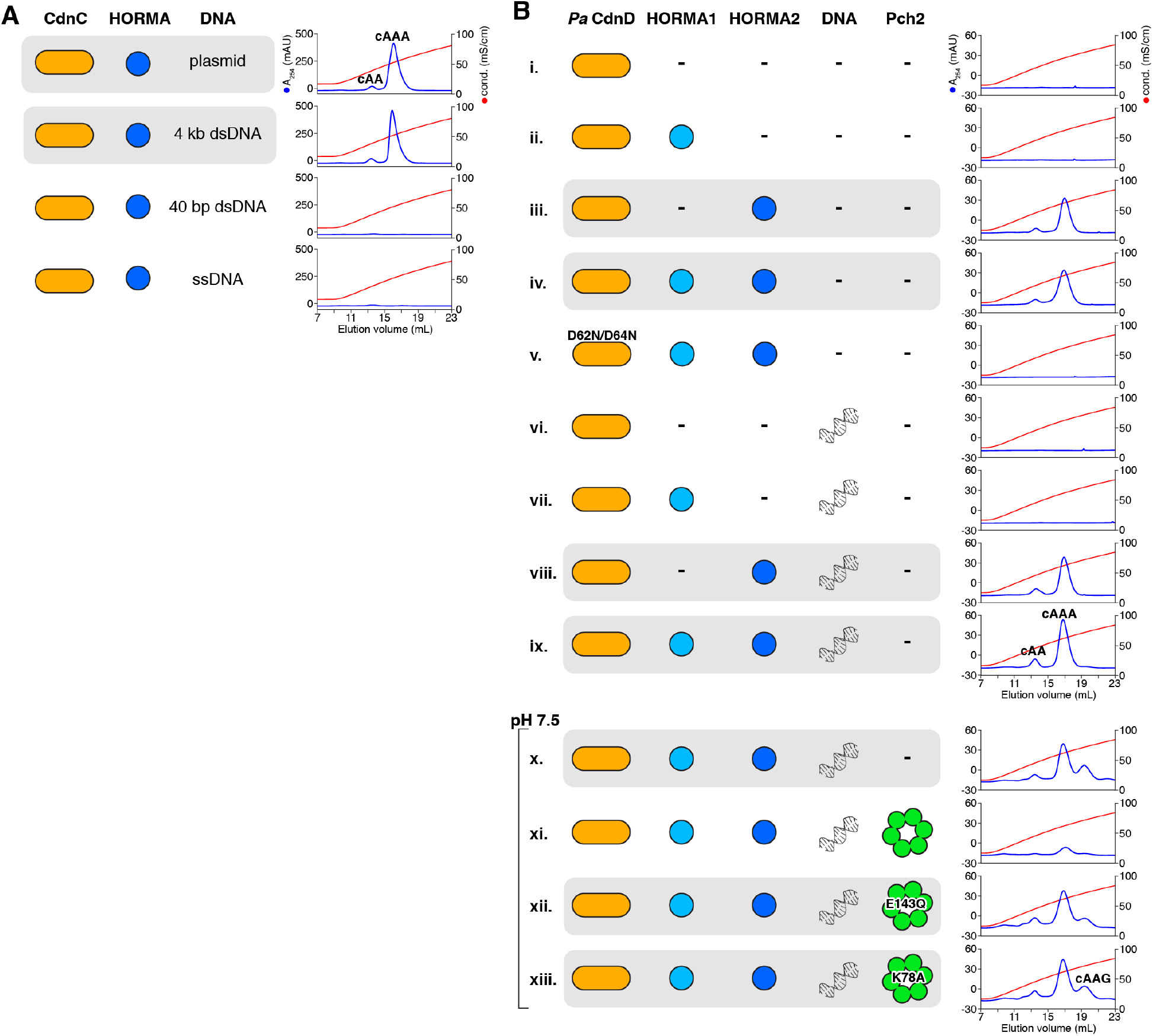
Second messenger synthesis by *Ec* CdnC and *Pa* CdnD. **(A)** Second messenger synthesis assays as in Fig. 1C, with different DNAs: ~4 kb plasmid DNA (top), 4 kb linear dsDNA, 40 bp linear dsDNA, and single-stranded DNA (sheared salmon-sperm DNA). **(B)** Second messenger synthesis assays with *Pa* CdnD, HORMA1, HORMA2, plasmid DNA, and Pch2. The peaks labeled “cAA” (sample ix), “cAAA” (sample ix), and “cAAG” (sample xiii) were verified by LC MS/MS. Samples x-xiii were assayed at pH 7.5 rather than 8.5 (as in samples i-ix) to accommodate *Pa* Pch2, which was insoluble at pH 8.5. *Pa* CdnD synthesizes cAAG as a minor product at pH 7.5.

**Figure S4.**
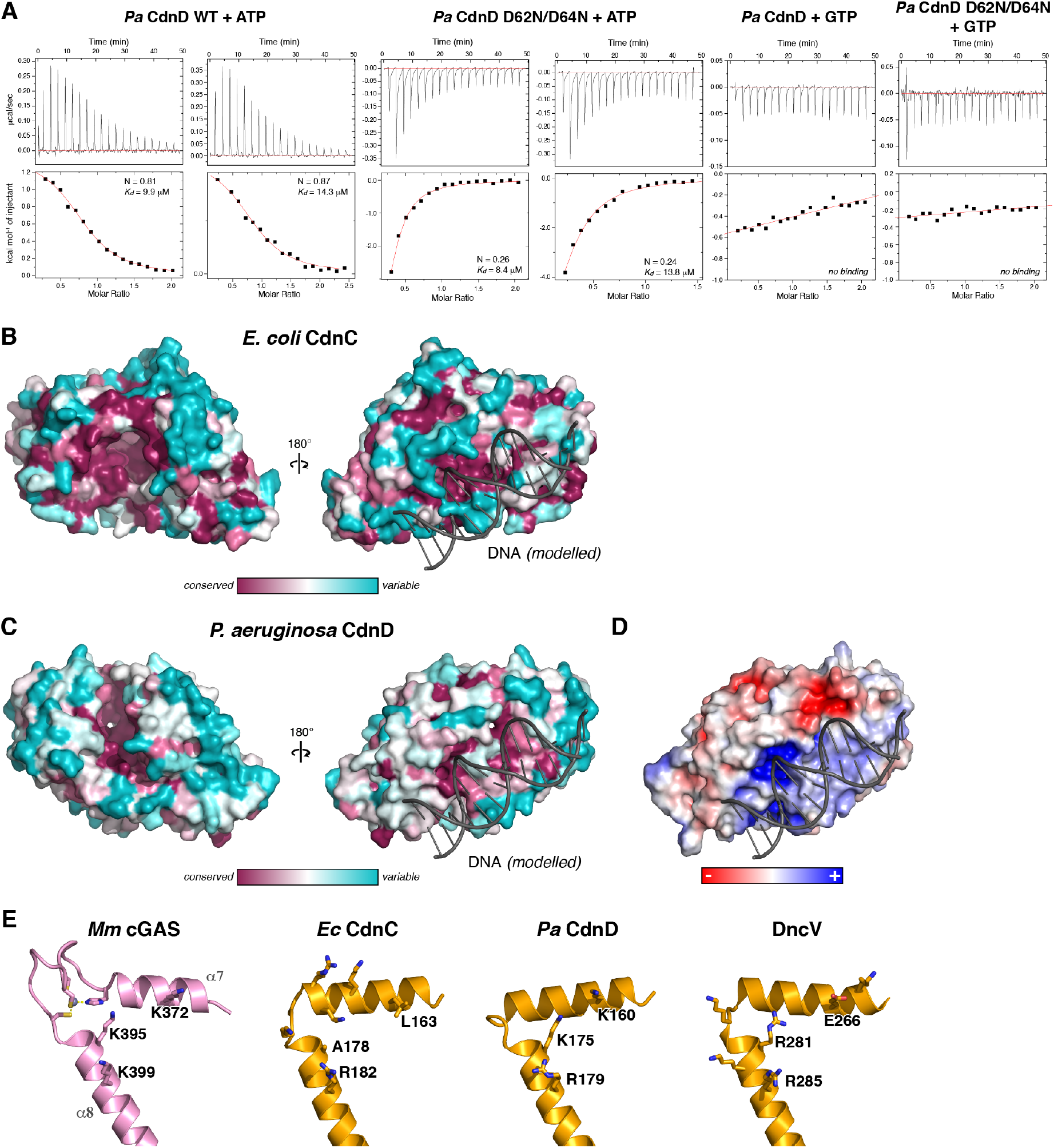
Structure and nucleotide binding of *Ec* CdnC and *Pa* CdnD. **(A)** Isothermal titration calorimetry (ITC) analysis of *Pa* CdnD binding either ATP or GTP. Both wild-type and D62N/D64N mutants bind ATP with a *K_d_* of ~ 10 μM, but neither detectably binds GTP. A similar analysis of *Ec* CdnC was not possible due to its stably-bound ATP. (B) Two views of *Ec* CdnC (left panel equivalent to Fig. 2A), with surface colored by conservation. Bound DNA (gray) is modelled from a superposition with *M. musculus* cGAS bound to DNA (PDB 4K9B) (Gao et al., 2013). **(C)** Two views of *Pa* CdnD equivalent (left panel equivalent to Fig. 2C, with surface colored by conservation. **(D)** View of *Pa* CdnD colored by surface charge. **(E)** Closeup view of the DNA-binding surface of *M. musculus* cGAS (PDB ID 4K9B) (Zhang et al. 2014), and equivalent regions of *Ec* CdnC, *Pa* CdnD, and *V. cholerae* DncV (PDB ID 4TXZ) (Kranzusch et al. 2014). cGAS residues K372, K395, and K399 (equivalent to human cGAS residues K384, K407, and K411) all contact DNA and are required for cGAS function (Kato et al. 2013; Li et al. 2013; Zhang et al. 2014). Structurally-equivalent residues of *Ec* CdnC, *Pa* CdnD, and *V. cholerae* DncV are shown as sticks, and labeled. Not labeled are additional surface-exposed positively-charged residues in each protein (*Ec* CdnC K167, K170, R171, K175; DncV K263, K278, K282).

**Figure S5.**
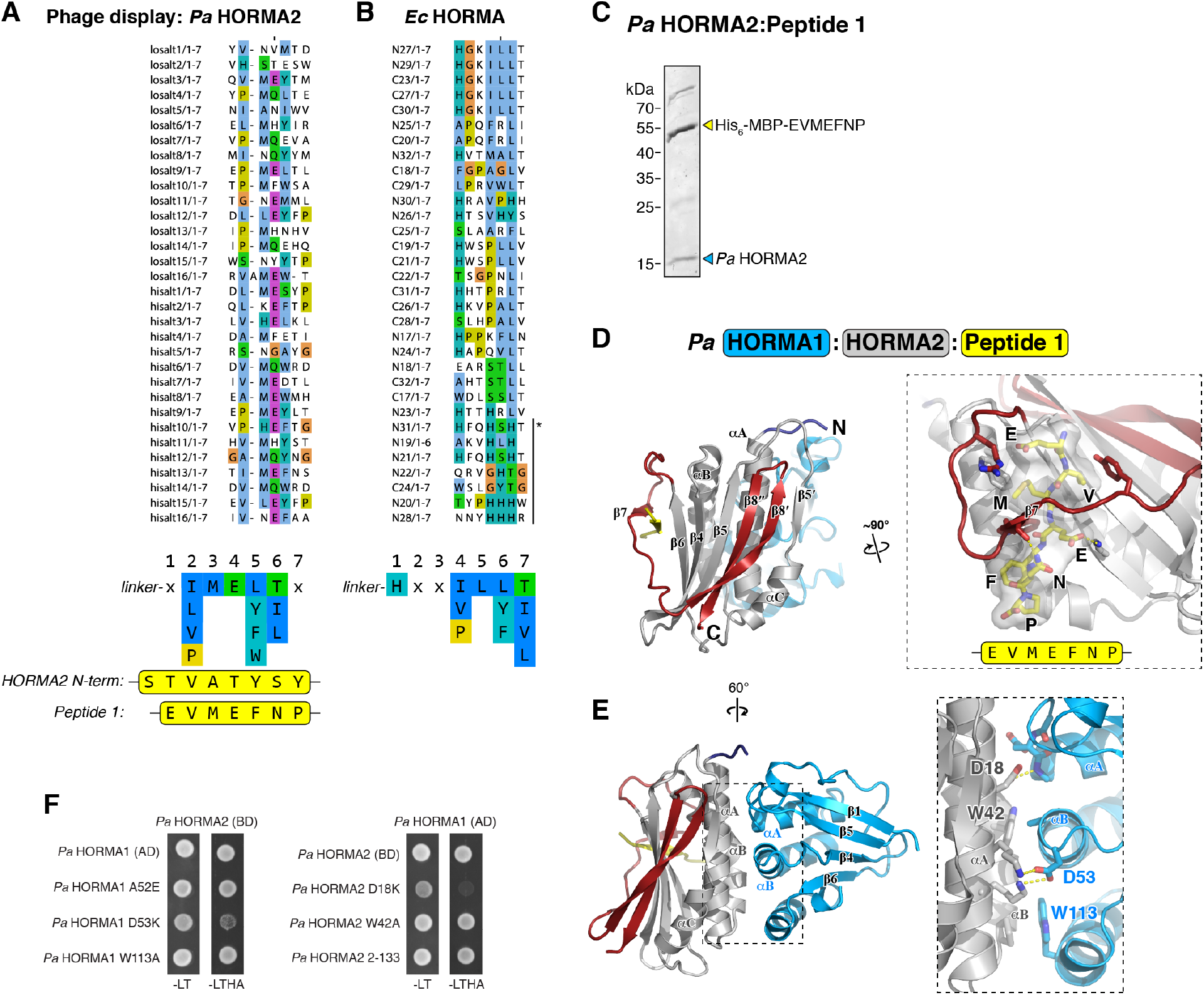
HORMA-closure motif and HORMA-HORMA interactions. **(A)** Aligned sequences from round 4 of phage display with *Pa* HORMA2 as bait. A consensus sequence is shown at bottom, aligned with the sequence of the *Pa* HORMA2 N-terminus, which binds *Pa* HORMA2 as a closure motif in our crystal structure (Fig. 3B), and Peptide 1, designed from the consensus motif. **(B)** Aligned sequences from round 4 of phage display with *Ec* HORMA as bait. A consensus sequence is shown at bottom. Asterisk indicated histidine-rich sequences, which may have bound non-specifically to the Ni-NTA resin used for selection. **(C)** Ni-NTA purification of coexpressed His6-MBP-Peptide **1** (sequence: EVMEFNP) with untagged *Pa* HORMA2. **(D)** Structure of the *Pa* HORMA1:HORMA2: Peptide 1 complex, with closeup view of HORMA2 binding Peptide 1 (yellow). **(E)** Rotated view of the *Pa* HORMA1:HORMA2:Peptide 1 complex, with closeup showing the HORMA1-HORMA2 interface. **(F)** Yeast two-hybrid analysis of the *Pa* HORMA1-HORMA2 interaction. BD: fusion to the Gal4 DNA-binding domain; AD: fusion to the Gal4 activation domain. -LT: media lacking leucine and tryptophan (non-selective); -LTHA: media lacking leucine, tryptophan, histidine, and adenine (stringent selection).

**Figure S6.**
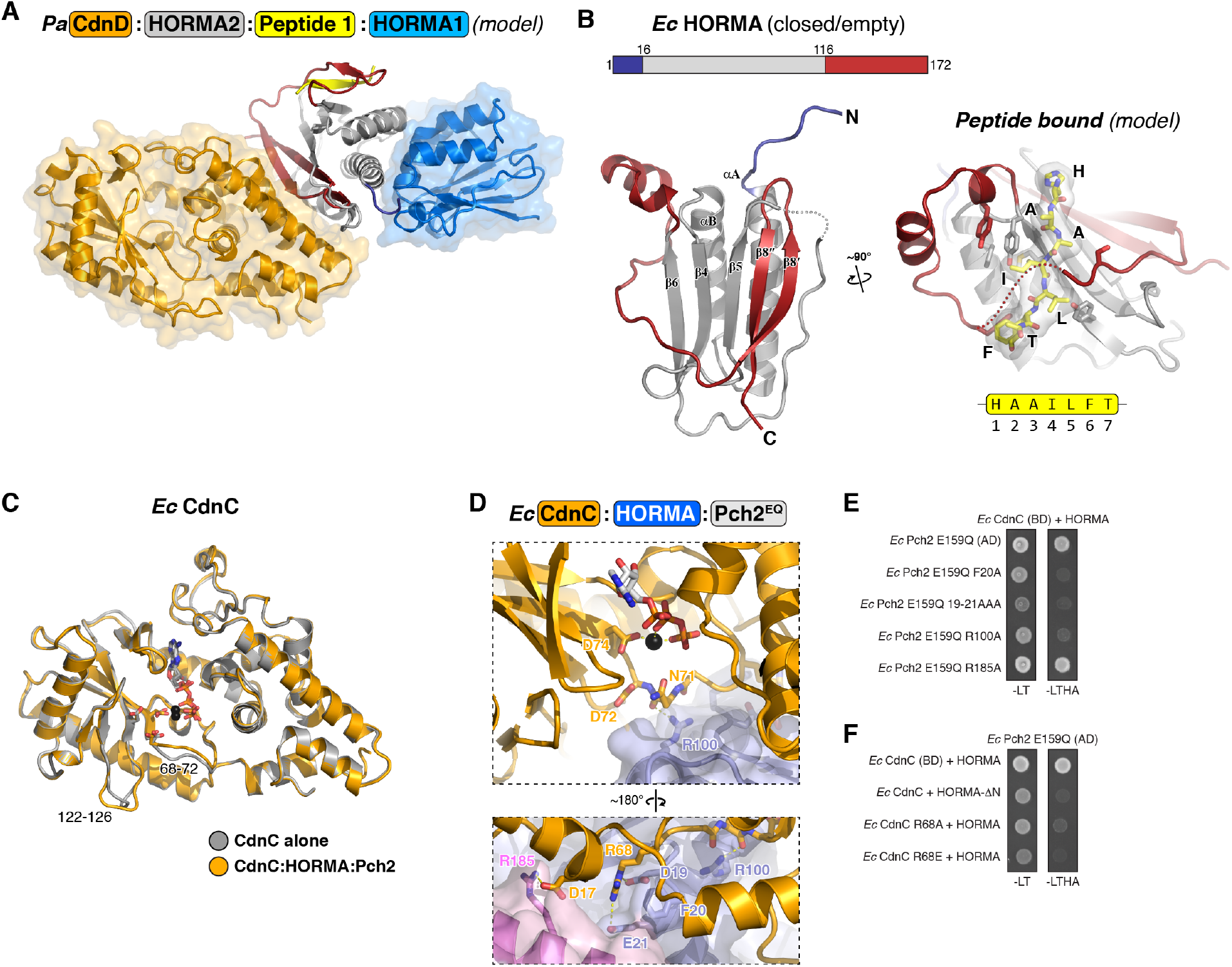
Structure of CD-NTase:HORMA complexes. **(A)** Model of a *Pa* CdnD:HORMA2:Peptide 1:HORMA1 complex, based on separate structures of CdnD:HORMA2:Peptide 1 (Fig. 4A) and HORMA1:HORMA2:Peptide 1 (Fig. 3E). HORMA1 and CdnD occupy different surfaces of HORMA2, suggesting they can bind simultaneously. **(B)** *Left:* Overall structure of *Ec* HORMA in the *Ec* CdnC:HORMA:Pch2 complex. Coloring is equivalent to Fig. 3A-D, with the mobile N-terminus blue and safety belt red. The protein adopts a closed, but empty, conformation. *Right:* Model of *Ec* HORMA bound to a consensus closure motif peptide (HAAILFT), based on a superposition of strands β5 and β6 of *Ec* HORMA with the *Pa* HORMA2:Peptide 1 complex. Residues 137-140 of the *Ec* HORMA safety belt, which clash with the bound peptide in the model, are shown as a dotted line. **(C)** Structural overlay of *Ec* CdnC alone (gray) with CdnC in complex with HORMA and Pch2 (orange). The overall Cα r.m.s.d. of the two structures is 0.3 Å; the most significant conformational changes are in residues 68-72 and 122-126, which contact Pch2. Despite directly contacting residues close to the CdnC active site (D72/D74), Pch2 binding does not significantly alter ATP Mg^2+^ binding. **(D)** Closeup views of the *Ec* CdnC-Pch2^EQ^ interface, with CdnC orange and Pch2^EQ^ monomers E and F shown as light blue and violet, respectively. *Top* shows closeup view equivalent to that of panel A, and *bottom* shows opposite view. **(E)** Yeast three-hybrid analysis (Ec Pch2 in pGADT7, *Ec* CdnC (AD fusion) and HORMA (untagged) in pBridge) showing that mutation of Pch2^EQ^ residues Asp19, Phe20, Glu21, and Arg100 disrupt binding to CdnC. -LT: media lacking leucine and tryptophan (non-selective); -LTHA: media lacking leucine, tryptophan, histidine, and adenine (stringent selection). **(F)** Yeast three-hybrid analysis showing that mutation of *Ec* CdnC residue Arg68, or truncation of the HORMA N-terminus, disrupt binding to Pch2^EQ^.

**Figure S7.**
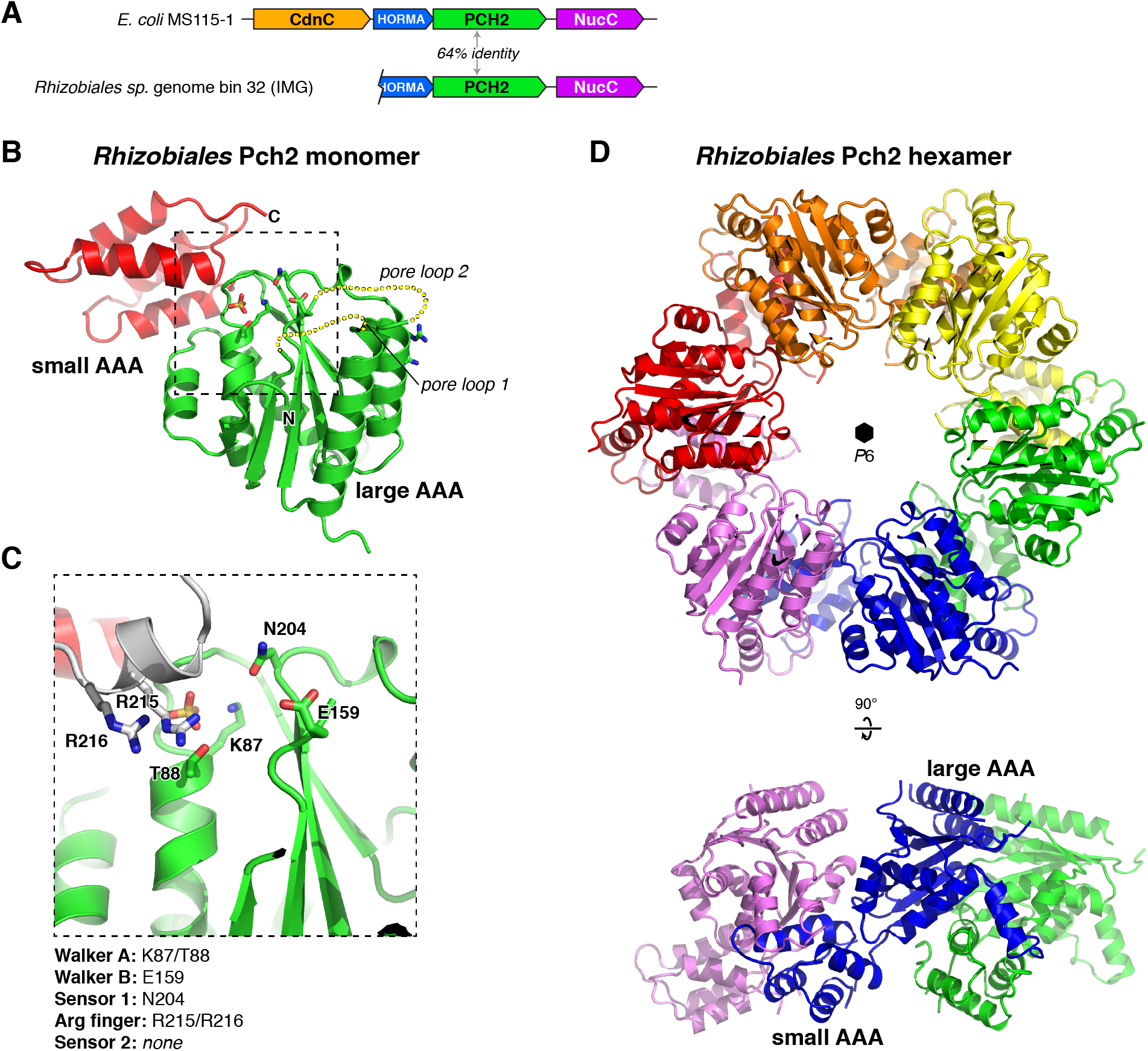
Structure of Rhizobiales Pch2. **(A)** Schematic of the *Rhizobiales* CD-NTase operon, including HORMA (IMG #2619783695), Pch2 (IMG #2619783694, sequence in Table S2), and NucC (IMG #2619783693), compared to the *E. coli* MS115-1 operon. The sequenced contig at IMG (JGI Integrated Microbial Genomes database) ends at the putative HORMA protein, precluding identification of the inferred CD-NTase. **(B)** Overall structure of the *Rhizobiales* (*Rh*) Pch2 monomer, with N-terminal “large AAA” domain green and C-terminal “small AAA” domain red. Yellow dotted lines indicate the positions of pore loops 1 and 2, both disordered in this structure. **(C)** Closeup view of the *Rh* Pch2 active site, in the same orientation as in panel B. A SO_4_^−^ ion is bound in the active site, surrounded by canonical Walker ATPase active site residues. Bacterial Pch2 proteins from 1-HORMA operons possess a conserved double Arg finger (R215/R216; from neighboring subunit, colored white); see Fig. S8F for sequences. **(D)** Top and side views of the six-fold symmetric *Rh* Pch2 hexamer, formed from crystallographic packing interactions (*P6* symmetry axis noted by black hexagon).

**Figure S8.**
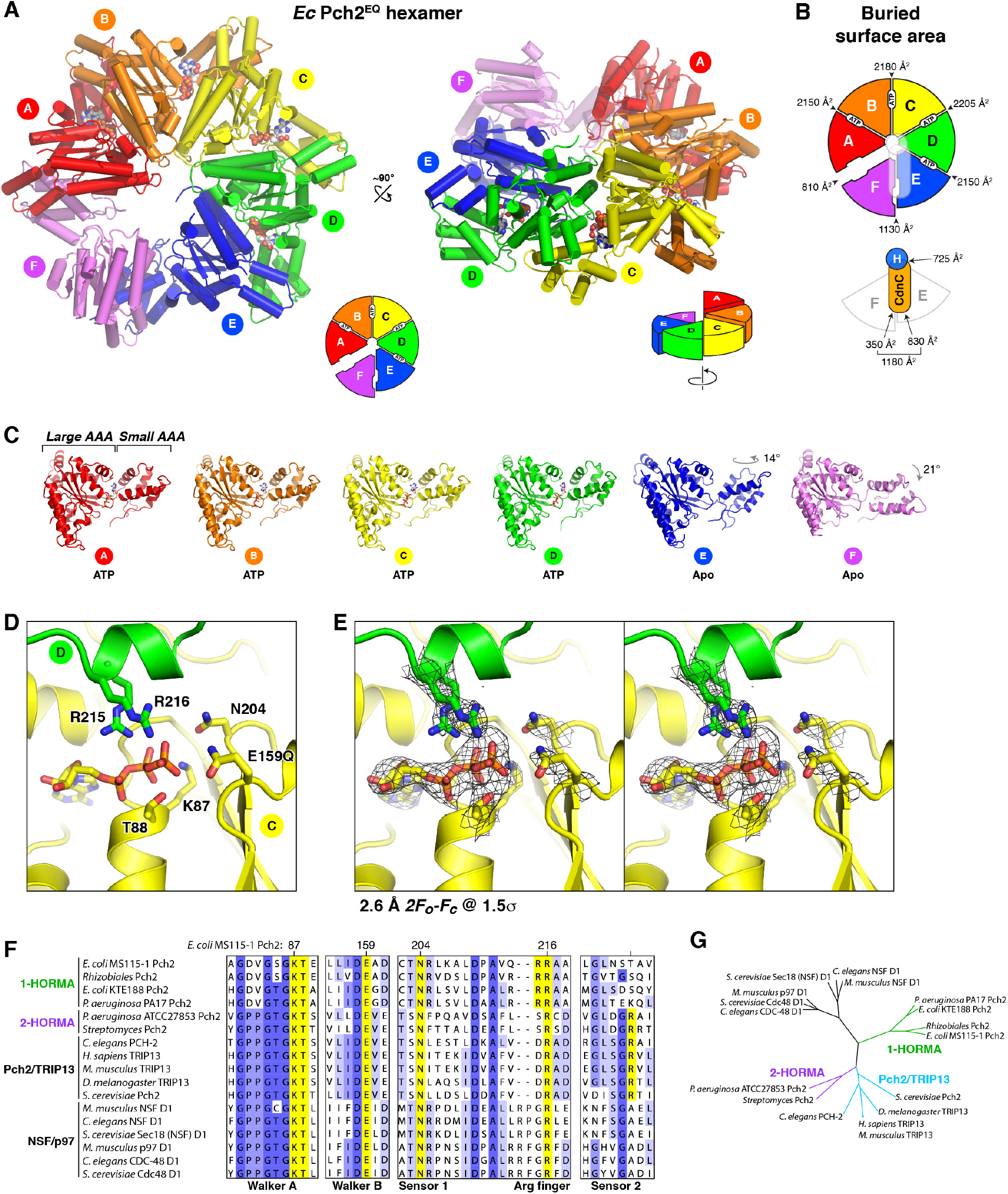
Structure of the *Ec* CdnC:HORMA:Pch2^EQ^ complex. **(A)** Top and side views of the *Ec* Pch2^EQ^ hexamer in the structure of CdnC:HORMA:Pch2^EQ^, with subunits A-F colored red, orange, yellow, green, blue, and violet, respectively. Bound ATP molecules at the A-B, B-C, C-D, and D-E interfaces are shown as spheres. **(B)** Schematics of the *Ec* CdnC:HORMA:Pch2^EQ^ complex, showing buried surface area in each subunit interface: inter-Pch2 interfaces top; Pch2-CdnC and CdnC-HORMA interfaces bottom. **(C)** Views of each individual subunit in the *Ec* Pch2^EQ^ hexamer, aligned on the large AAA domain, showing the conformational similarity of subunits A-D, and rotation of the small AAA domain in subunits E and F. **(D)** Closeup of ATP binding to *Ec* Pch2^EQ^ (subunit C), with conserved ATP-interacting residues from chain C (Walker A, Walker B, Sensor 1) in yellow and from chain D (double Arg finger) in green. **(E)** Stereo view of refined 2*F_o_-F_c_* electron density, contoured at 1.5 σ, surrounding ATP and ATP-interacting residues in Pch2^EQ^ chain C. **(F)** Conserved AAA+ sequence motifs in bacterial Pch2, eukaryotic Pch2/TRIP13, and eukaryotic NSF and p97 proteins (D1 ATPase domain). While Pch2 proteins from 2-HORMA operons (e.g. *P. aeruginosa* ATCC27853 Pch2) share all motifs with eukaryotic Pch2/TRIP13 proteins, Pch2 proteins from 1-HORMA operons (e.g. *E. coli* MS115-1 Pch2) possess a double Arg finger (*Ec* Pch2 residues 215 and 216) and lack a sensor 2 arginine. **(G)** Phylogenetic tree showing that bacterial Pch2 proteins from 2-HORMA operons are more closely related to eukaryotic Pch2/TRIP13 than those from 1-HORMA operons.

**Figure S9.**
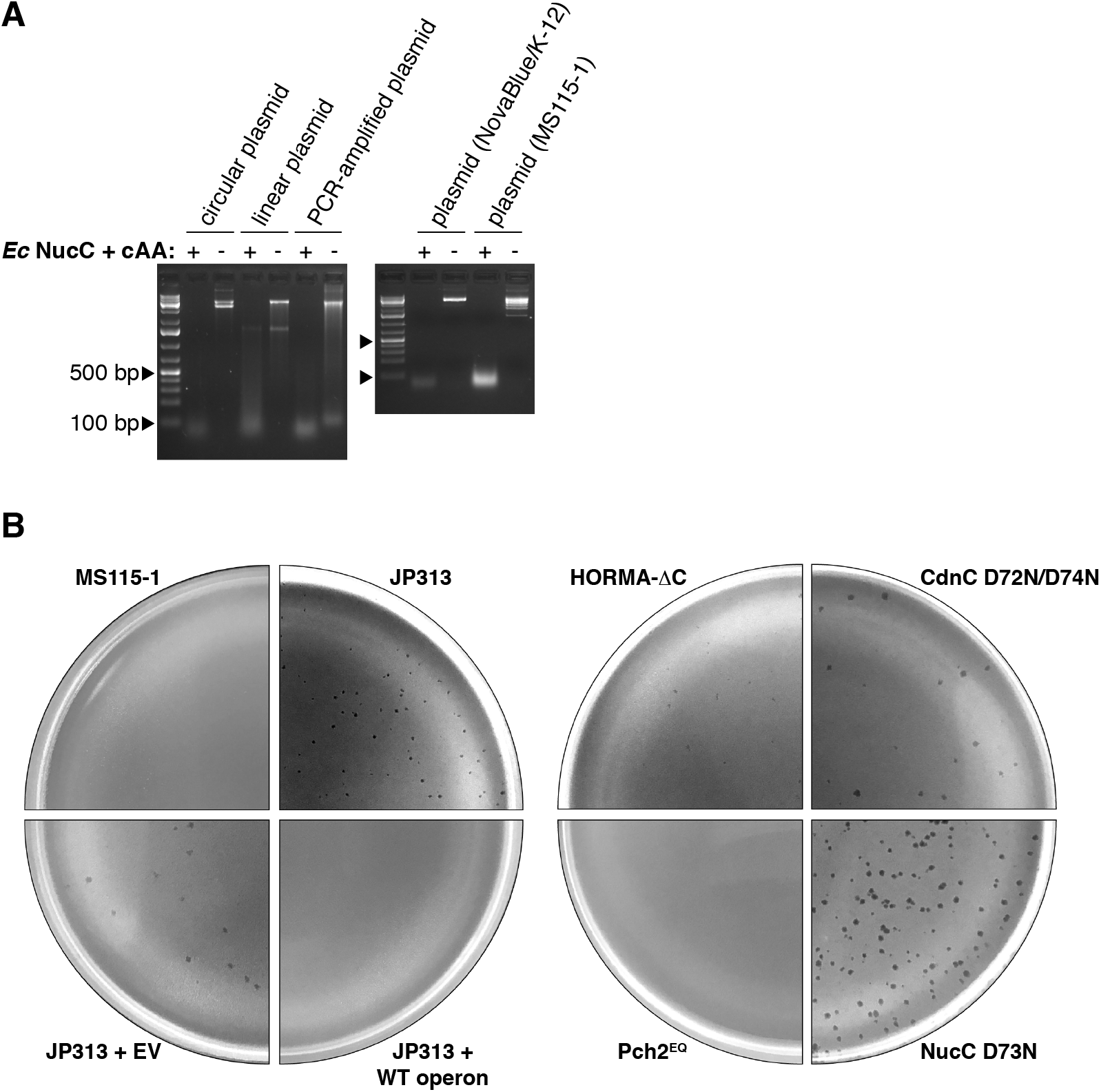
NucC activity and bacteriophage immunity. **(A)** Plasmid digestion assay with 10 nM *Ec* NucC, 1 μM cAA, and 1 μg of the indicated DNAs. **(B)** Representative images of plates used for counting bacteriophage plaques (Fig. 6C).

## Supplemental Note

Sequence of *E. coli* MS115-1 CD-NTase operon cloned into plasmid pLAC22 for plaque assays:

Color-coding:

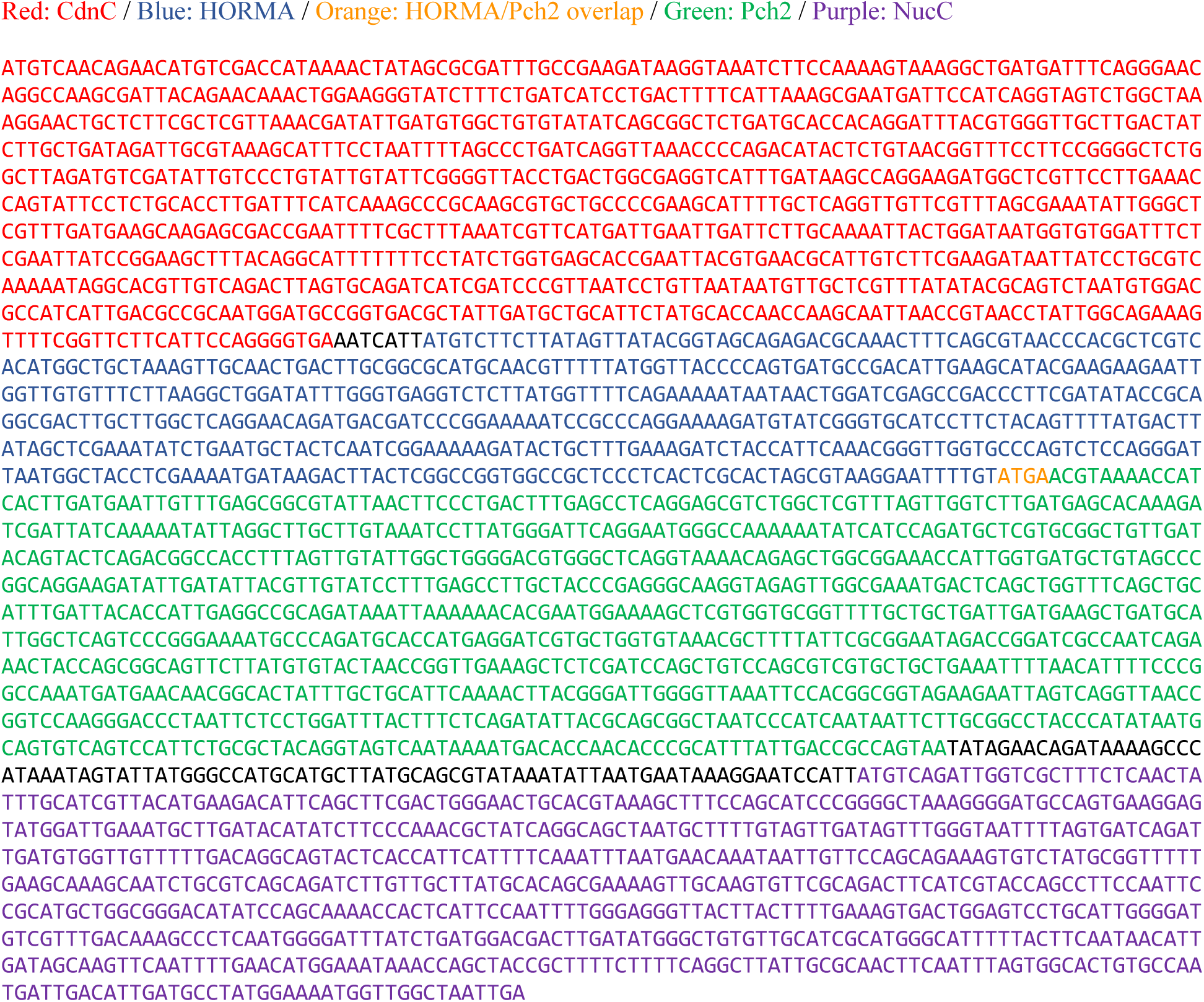

**Table S1 –.**
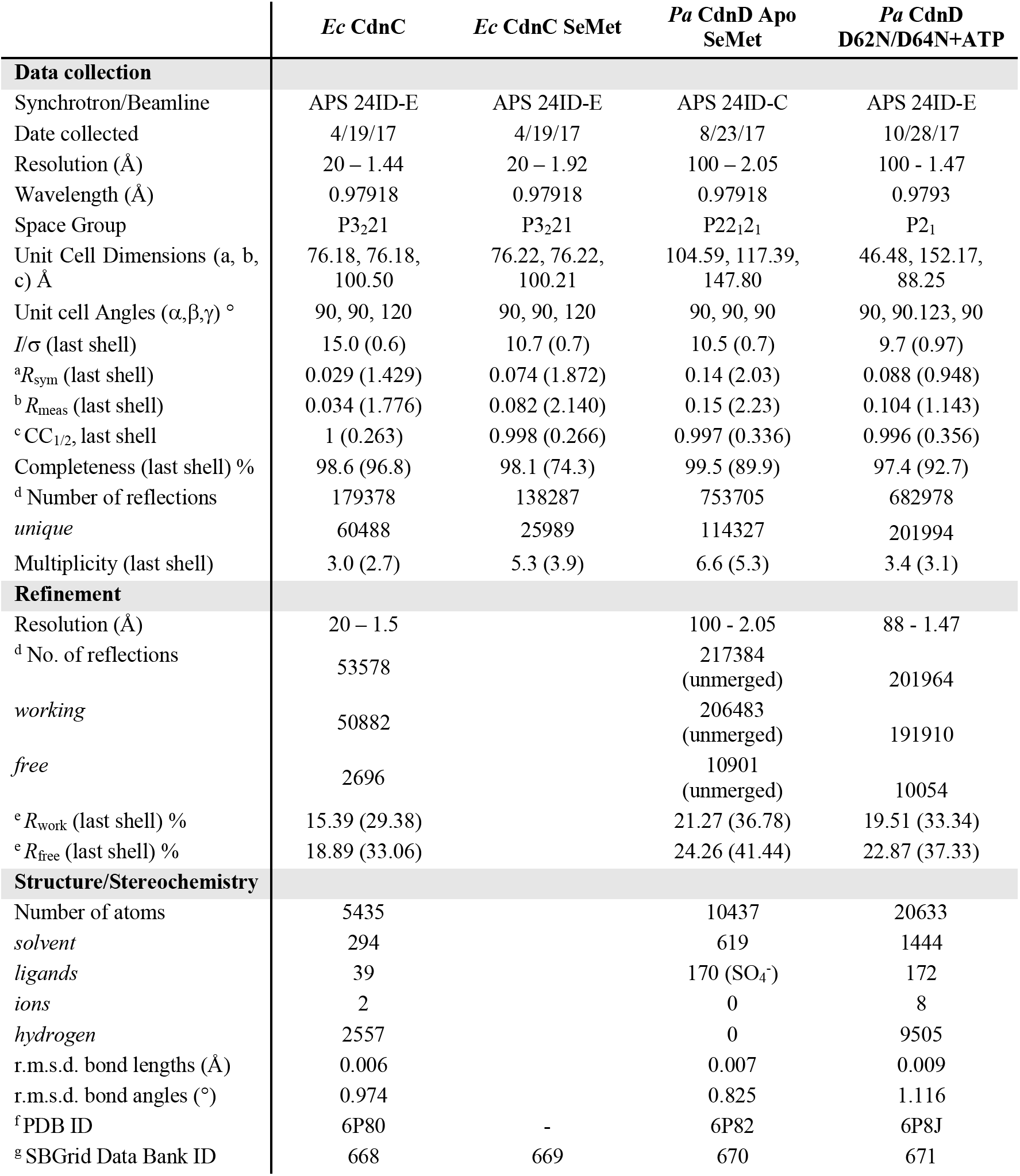
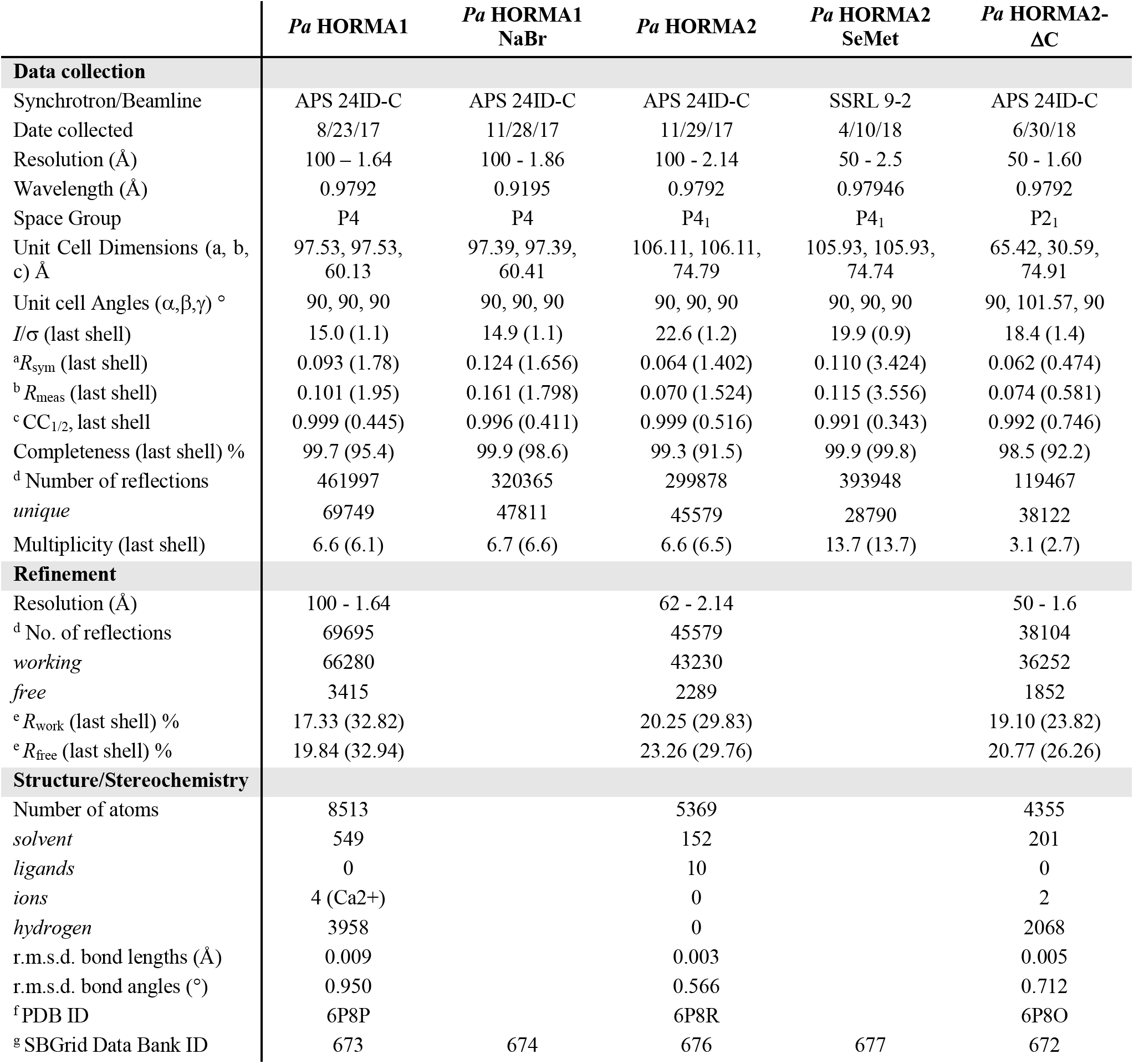
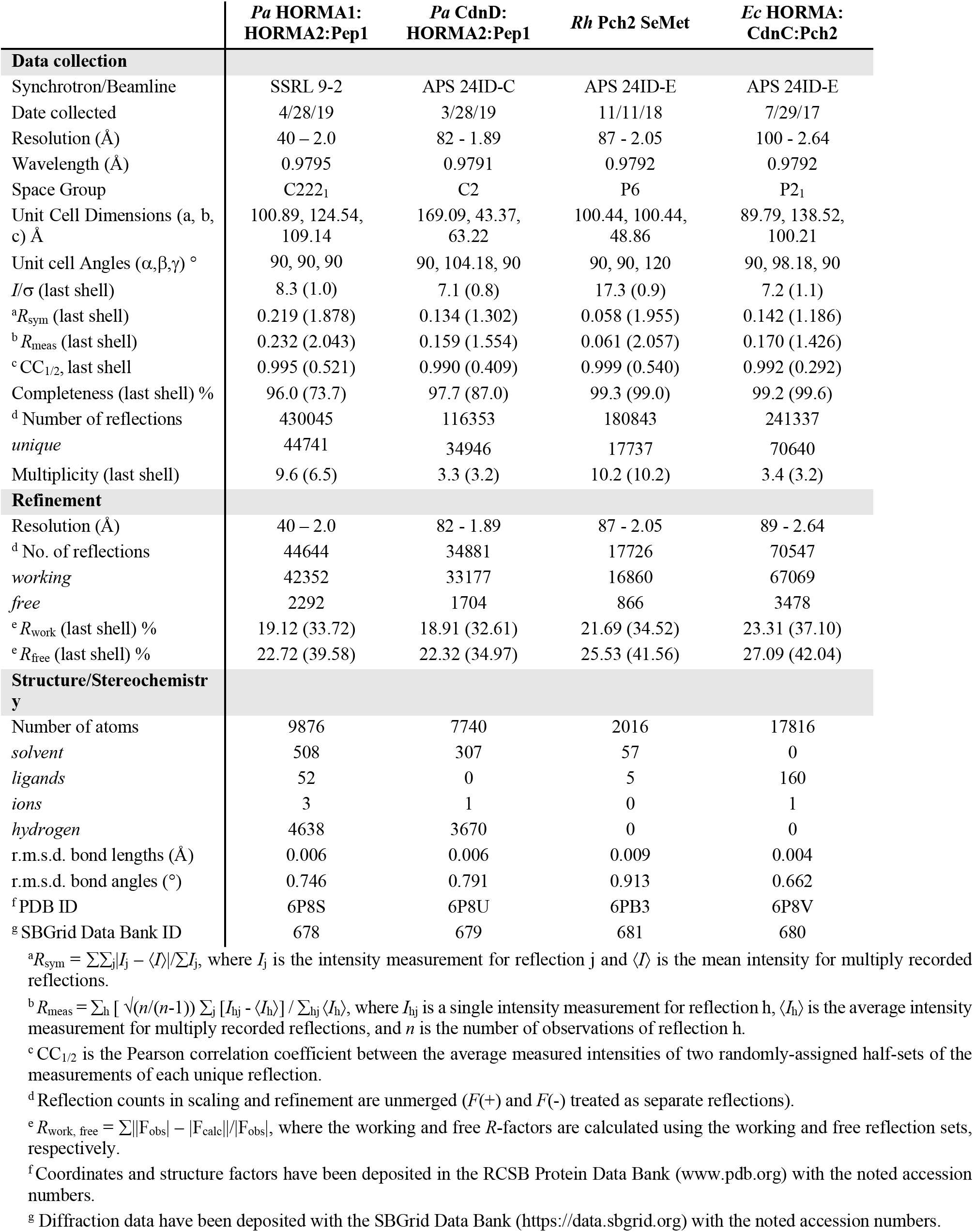
Crystallographic data collection and refinement

**Table S2 –.**
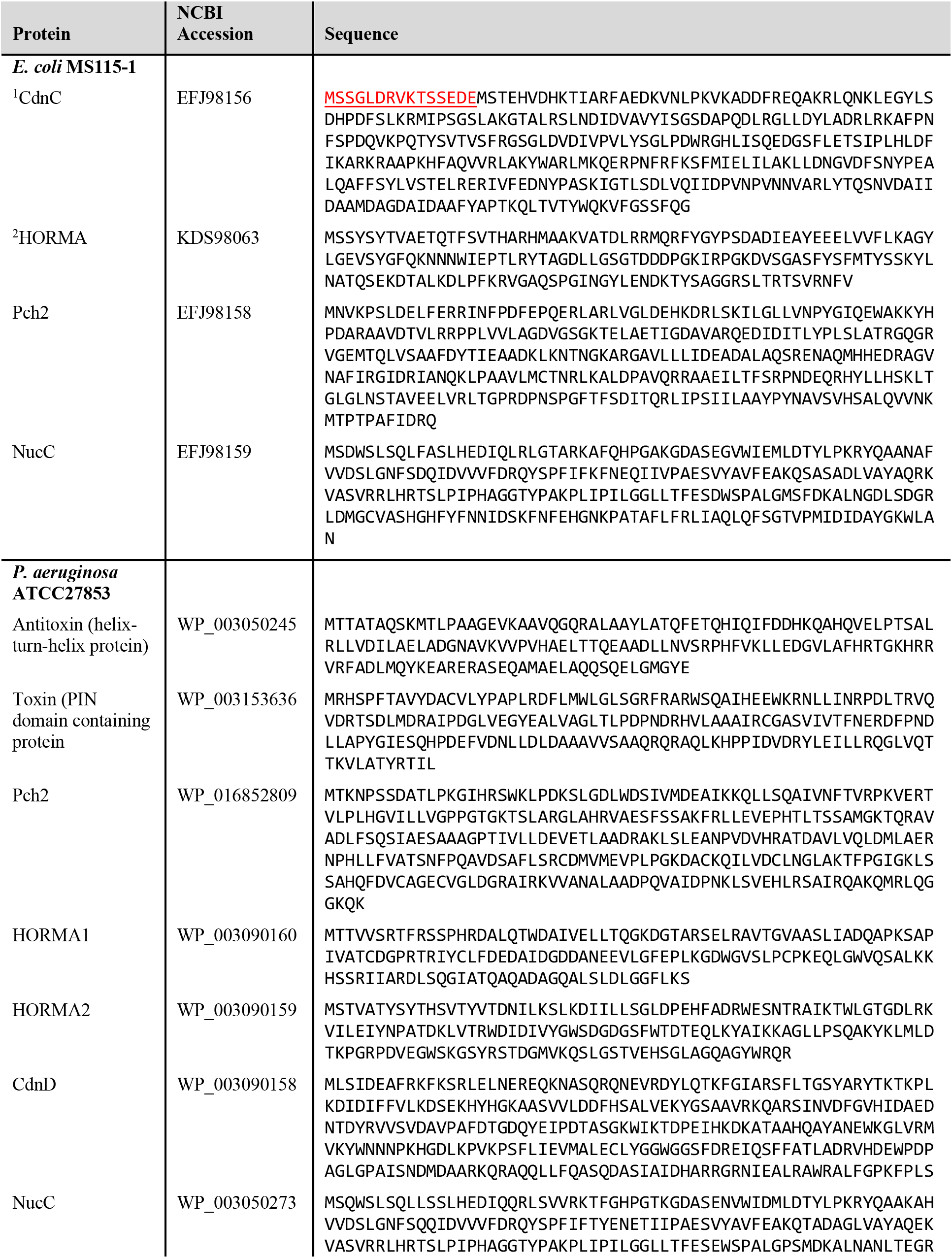
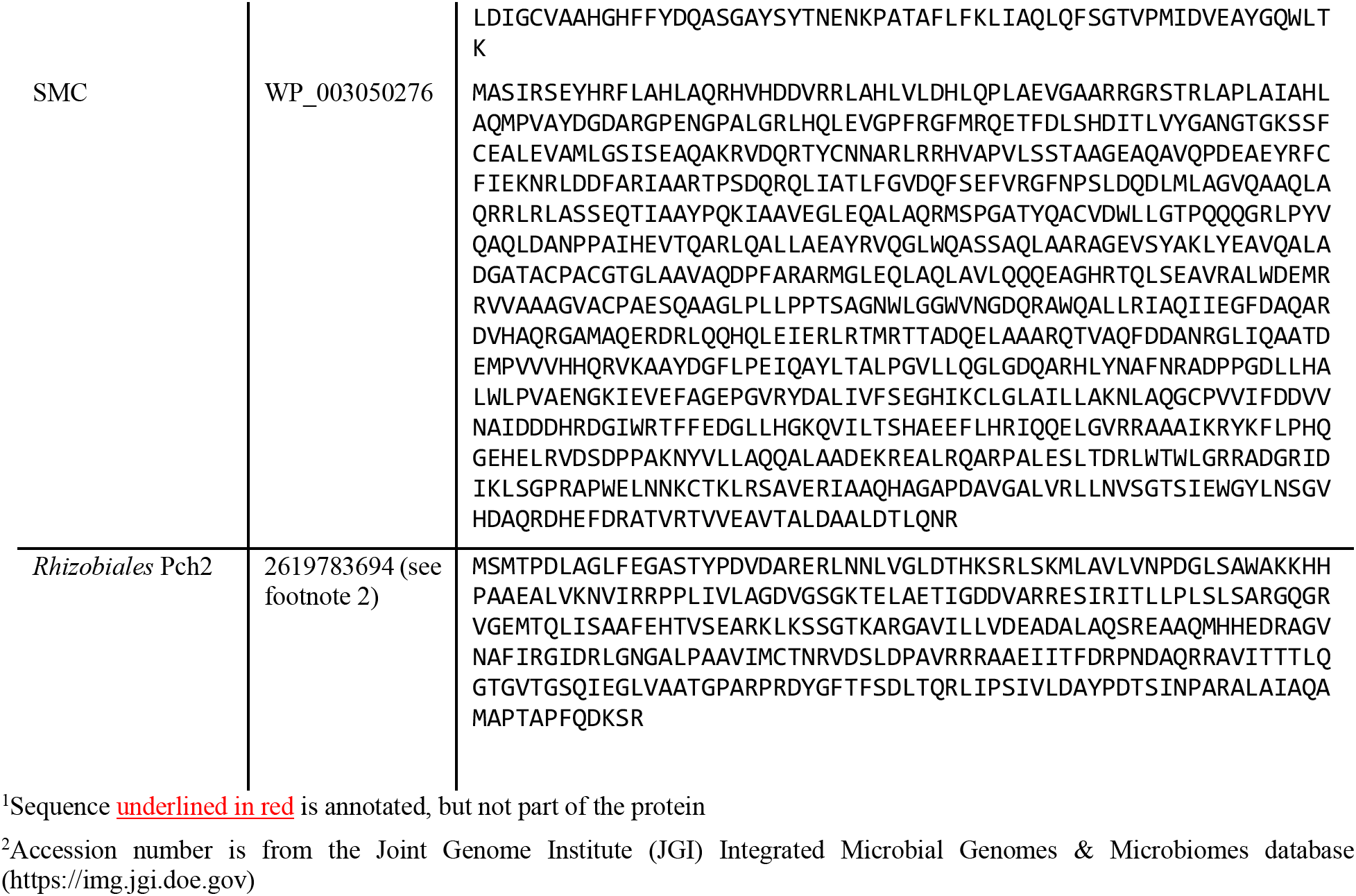
Protein Sequences

## References

Afonine, P. V, Grosse-Kunstleve, R.W., Echols, N., Headd, J.J., Moriarty, N.W., Mustyakimov, M., Terwilliger, T.C., Urzhumtsev, A., Zwart, P.H., Adams, P.D., et al. (2012). Towards automated crystallographic structure refinement with phenix.refine. Acta Crystallogr. Sect. D, Biol. Crystallogr. 68, 352–367.

Alfieri, C., Chang, L., and Barford, D. (2018). Mechanism for remodelling of the cell cycle checkpoint protein MAD2 by the ATPase TRIP13. Nature 559, 274–278.

Aravind, L., and Koonin, E. V (1998). The HORMA domain: a common structural denominator in mitotic checkpoints, chromosome synapsis and DNA repair. Trends Biochem. Sci. 23, 284–286.

Burdette, D.L., Monroe, K.M., Sotelo-Troha, K., Iwig, J.S., Eckert, B., Hyodo, M., Hayakawa, Y., and Vance, R.E. (2011). STING is a direct innate immune sensor of cyclic di-GMP. Nature 478, 515–518.

Burroughs, A.M., Zhang, D., Schäffer, D.E., Iyer, L.M., and Aravind, L. (2015). Comparative genomic analyses reveal a vast, novel network of nucleotide-centric systems in biological conflicts, immunity and signaling. Nucleic Acids Res. 43, gkv1267–10654.

Chen, Q., Sun, L., and Chen, Z.J. (2016). Regulation and function of the cGAS–STING pathway of cytosolic DNA sensing. Nat. Immunol. 17, 1142–1149.

Civril, F., Deimling, T., de Oliveira Mann, C.C., Ablasser, A., Moldt, M., Witte, G., Hornung, V., and Hopfner, K.-P. (2013). Structural mechanism of cytosolic DNA sensing by cGAS. Nature 498, 332–337.

Corrigan, R.M., and Gründling, A. (2013). Cyclic di-AMP: another second messenger enters the fray. Nat. Rev. Microbiol. 11, 513–524.

Davies, B.W., Bogard, R.W., Young, T.S., and Mekalanos, J.J. (2012). Coordinated regulation of accessory genetic elements produces cyclic dinucleotides for V. cholerae virulence. Cell 149, 358–370.

Deville, C., Carroni, M., Franke, K.B., Topf, M., Bukau, B., Mogk, A., and Saibil, H.R. (2017). Structural pathway of regulated substrate transfer and threading through an Hsp100 disaggregase. Sci. Adv. 3, e1701726.

Van Duyne, G.D., Standaert, R.F., Karplus, P.A., Schreiber, S.L., and Clardy, J. (1993). Atomic structures of the human immunophilin FKBP-12 complexes with FK506 and rapamycin. J. Mol. Biol. 229, 105–124.

Economou, A., Pogliano, J.A., Beckwith, J., Oliver, D.B., and Wickner, W. (1995). SecA membrane cycling at SecYEG is driven by distinct ATP binding and hydrolysis events and is regulated by SecD and SecF. Cell 83, 1171–1181.

Emsley, P., Lohkamp, B., Scott, W.G., and Cowtan, K. (2010). Features and development of Coot. Acta Crystallogr. Sect. D, Biol. Crystallogr. 66, 486–501.

Evans, P.R., and Murshudov, G.N. (2013). How good are my data and what is the resolution? Acta Crystallogr. Sect. D, Biol. Crystallogr. 69, 1204–1214.

Gao, P., Ascano, M., Wu, Y., Barchet, W., Gaffney, B.L., Zillinger, T., Serganov, A.A., Liu, Y., Jones, R.A., Hartmann, G., et al. (2013). Cyclic [G(2’,5’)pA(3’,5’)p] is the metazoan second messenger produced by DNA-activated cyclic GMP-AMP synthase. Cell 153, 1094–1107.

Gates, S.N., Yokom, A.L., Lin, J., Jackrel, M.E., Rizo, A.N., Kendsersky, N.M., Buell, C.E., Sweeny, E.A., Mack, K.L., Chuang, E., et al. (2017). Ratchet-like polypeptide translocation mechanism of the AAA+ disaggregase Hsp104. Sci. (New York, NY) 357, 273–279.

Han, H., Monroe, N., Sundquist, W.I., Shen, P.S., and Hill, C.P. (2017). The AAA ATPase Vps4 binds ESCRT-III substrates through a repeating array of dipeptide-binding pockets. Elife 6, 50.

Hengge, R. (2009). Principles of c-di-GMP signalling in bacteria. Nat. Rev. Microbiol. 7, 263–273.

van Hooff, J.J., Tromer, E., van Wijk, L.M., Snel, B., and Kops, G.J. (2017). Evolutionary dynamics of the kinetochore network in eukaryotes as revealed by comparative genomics. EMBO Rep. 18, 1559–1571.

Hornung, V., Hartmann, R., Ablasser, A., and Hopfner, K.-P. (2014). OAS proteins and cGAS: unifying concepts in sensing and responding to cytosolic nucleic acids. Nat. Rev. Immunol. 14, 521–528.

Jones, D.T., and Cozzetto, D. (2015). DISOPRED3: precise disordered region predictions with annotated protein-binding activity. Bioinformatics 31, 857–863.

Kabsch, W. (2010). XDS. Acta Crystallogr. Sect. D, Biol. Crystallogr. 66, 125–132.

Kato, K., Ishii, R., Goto, E., Ishitani, R., Tokunaga, F., and Nureki, O. (2013). Structural and functional analyses of DNA-sensing and immune activation by human cGAS. PLoS One 8, e76983.

Kim, D.H., Han, J.S., Ly, P., Ye, Q., McMahon, M.A., Myung, K., Corbett, K.D., and Cleveland, D.W. (2018). TRIP13 and APC15 drive mitotic exit by turnover of interphase- and unattached kinetochore-produced MCC. Nat. Commun. 9, 4354.

Lau, R.K., Ye, Q., Berg, K.R., Mathews, I.T., Watrous, J.D., Whiteley, A.T., Lowey, B., Mekalanos J.J., Kranzusch, P.J., Jain, M., and Corbett, K.D. (2019). Structure and mechanism of a cyclic trinucleotide-activated bacterial endonuclease mediating bacteriophage immunity. bioRxiv 694703. https://doi.org/10.1101/694703

Li, X., Shu, C., Yi, G., Chaton, C.T., Shelton, C.L., Diao, J., Zuo, X., Kao, C.C., Herr, A.B., and Li, P. (2013). Cyclic GMP-AMP synthase is activated by double-stranded DNA-induced oligomerization. Immunity 39, 1019–1031.

Livingstone, C.D., and Barton, G.J. (1993). Protein sequence alignments: a strategy for the hierarchical analysis of residue conservation. Comput. Appl. Biosci. 9, 745–756.

Lohöfener, J., Steinke, N., Kay-Fedorov, P., Baruch, P., Nikulin, A., Tishchenko, S., Manstein, D.J., and Fedorov, R. (2015). The Activation Mechanism of 2’-5’-Oligoadenylate Synthetase Gives New Insights Into OAS/cGAS Triggers of Innate Immunity. Structure 23, 851–862.

Luo, X., and Yu, H. (2008). Protein metamorphosis: the two-state behavior of Mad2. Structure 16, 1616–1625.

Ma, H.T., and Poon, R.Y.C. (2016). TRIP13 Regulates Both the Activation and Inactivation of the Spindle-Assembly Checkpoint. Cell Rep. 14, 1086–1099.

Ma, H.T., and Poon, R.Y.C. (2018). TRIP13 Functions in the Establishment of the Spindle Assembly Checkpoint by Replenishing O-MAD2. Cell Rep. 22, 1439–1450.

Mapelli, M., Massimiliano, L., Santaguida, S., and Musacchio, A. (2007). The Mad2 Conformational Dimer: Structure and Implications for the Spindle Assembly Checkpoint. Cell 131, 730–743.

McCoy, A.J., Grosse-Kunstleve, R.W., Adams, P.D., Winn, M.D., Storoni, L.C., and Read, R.J. (2007). Phaser crystallographic software. J. Appl. Crystallogr. 40, 658–674.

McFarland, A.P., Luo, S., Ahmed-Qadri, F., Zuck, M., Thayer, E.F., Goo, Y.A., Hybiske, K., Tong, L., and Woodward, J.J. (2017). Sensing of Bacterial Cyclic Dinucleotides by the Oxidoreductase RECON Promotes NF-κB Activation and Shapes a Proinflammatory Antibacterial State. Immunity 46, 433–445.

Pape, T., Schneider, T.R., and IUCr (2004). HKL2MAP: a graphical user interface for macromolecular phasing with SHELX programs. J. Appl. Crystallogr. 37, 843–844.

Puchades, C., Rampello, A.J., Shin, M., Giuliano, C.J., Wiseman, R.L., Glynn, S.E., and Lander, G.C. (2017). Structure of the mitochondrial inner membrane AAA+ protease YME1 gives insight into substrate processing. Sci. (New York, NY) 358, eaao0464.

Qi, S., Kim, D.J., Stjepanovic, G., and Hurley, J.H. (2015). Structure of the Human Atg13-Atg101 HORMA Heterodimer: an Interaction Hub within the ULK1 Complex. Structure 23, 1848–1857.

Ripstein, Z.A., Huang, R., Augustyniak, R., Kay, L.E., and Rubinstein, J.L. (2017). Structure of a AAA+ unfoldase in the process of unfolding substrate. Elife 6, 43.

Rosenberg, S.C., and Corbett, K.D. (2015). The multifaceted roles of the HORMA domain in cellular signaling. J. Cell Biol. 211, 745–755.

Sheldrick, G.M. (2010). Experimental phasing with SHELXC/D/E: combining chain tracing with density modification. Acta Crystallogr. Sect. D, Biol. Crystallogr. 66, 479–485.

Suzuki, H., Kaizuka, T., Mizushima, N., and Noda, N.N. (2015). Structure of the Atg101–Atg13 complex reveals essential roles of Atg101 in autophagy initiation. Nat. Struct. Mol. Biol. 22, 572–580.

Terwilliger, T.C., Adams, P.D., Read, R.J., McCoy, A.J., Moriarty, N.W., Grosse-Kunstleve, R.W., Afonine, P. V, Zwart, P.H., and Hung, L.-W. (2009). Decision-making in structure solution using Bayesian estimates of map quality: the PHENIX AutoSol wizard. Acta Crystallogr. Sect. D, Biol. Crystallogr. 65, 582–601.

White, K.I., Zhao, M., Choi, U.B., Pfuetzner, R.A., and Brunger, A.T. (2018). Structural principles of SNARE complex recognition by the AAA+ protein NSF. Elife 7, 213.

Whiteley, A.T., Eaglesham, J.B., de Oliveira Mann, C.C., Morehouse, B.R., Lowey, B., Nieminen, E.A., Danilchanka, O., King, D.S., Lee, A.S.Y., Mekalanos, J.J., et al. (2019). Bacterial cGAS-like enzymes synthesize diverse nucleotide signals. Nature 567, 194–199.

Winn, M.D., Ballard, C.C., Cowtan, K.D., Dodson, E.J., Emsley, P., Evans, P.R., Keegan, R.M., Krissinel, E.B., Leslie, A.G.W., McCoy, A., et al. (2011). Overview of the CCP4 suite and current developments. Acta Crystallogr. Sect. D, Biol. Crystallogr. 67, 235–242.

Ye, Q., Rosenberg, S.C., Moeller, A., Speir, J.A., Su, T.Y., and Corbett, K.D. (2015). TRIP13 is a protein-remodeling AAA+ ATPase that catalyzes MAD2 conformation switching. Elife 2015, 1–44.

Ye, Q., Kim, D.H., Dereli, I., Rosenberg, S.C., Hagemann, G., Herzog, F., Tóth, A., Cleveland, D.W., and Corbett, K.D. (2017). The AAA+ ATPase TRIP13 remodels HORMA domains through N-terminal engagement and unfolding. EMBO J. 36, 2419–2434.

Yu, H., Lupoli, T.J., Kovach, A., Meng, X., Zhao, G., Nathan, C.F., and Li, H. (2018). ATP hydrolysis-coupled peptide translocation mechanism of Mycobacterium tuberculosis ClpB. Proc. Natl. Acad. Sci. 115, E9560–E9569.

Zhang, X., Wu, J., Du, F., Xu, H., Sun, L., Chen, Z., Brautigam, C.A., Zhang, X., and Chen, Z.J. (2014). The cytosolic DNA sensor cGAS forms an oligomeric complex with DNA and undergoes switch-like conformational changes in the activation loop. Cell Rep. 6, 421–430.

Zhu, D., Wang, L., Shang, G., Liu, X., Zhu, J., Lu, D., Wang, L., Kan, B., Zhang, J.-R., and Xiang, Y. (2014). Structural biochemistry of a Vibrio cholerae dinucleotide cyclase reveals cyclase activity regulation by folates. Mol. Cell 55, 931–937.

## Supplemental References

Gao P, Ascano M, Wu Y, et al (2013) Cyclic [G(2’,5’)pA(3’,5’)p] is the metazoan second messenger produced by DNA-activated cyclic GMP-AMP synthase. Cell 153:1094–1107. doi: 10.1016/j.cell.2013.04.046

Kato K, Ishii R, Goto E, et al (2013) Structural and functional analyses of DNA-sensing and immune activation by human cGAS. PLoS One 8:e76983. doi: 10.1371/journal.pone.0076983

Kranzusch PJ, Lee ASY, Wilson SC, et al (2014) Structure-guided reprogramming of human cGAS dinucleotide linkage specificity. Cell 158:1011–1021. doi: 10.1016/j.cell.2014.07.028

Li X, Shu C, Yi G, et al (2013) Cyclic GMP-AMP synthase is activated by double-stranded DNA-induced oligomerization. Immunity 39:1019–1031. doi: 10.1016/j.immuni.2013.10.019

Makarova KS, Wolf YI, Koonin E V (2009) Comprehensive comparative-genomic analysis of type 2 toxin-antitoxin systems and related mobile stress response systems in prokaryotes. Biol Direct 4:19. doi: 10.1186/1745-6150-4-19

Skjerning RB, Senissar M, Winther KS, et al (2018) The RES domain toxins of RES-Xre toxin-antitoxin modules induce cell stasis by degrading NAD. Mol Microbiol 66:213. doi: 10.1111/mmi.14150

Whiteley AT, Eaglesham JB, de Oliveira Mann CC, et al (2019) Bacterial cGAS-like enzymes synthesize diverse nucleotide signals. Nature 567:194–199. doi: 10.1038/s41586-019-0953-5

Zhang X, Wu J, Du F, et al (2014) The cytosolic DNA sensor cGAS forms an oligomeric complex with DNA and undergoes switch-like conformational changes in the activation loop. Cell Rep 6:421–30. doi: 10.1016/j.celrep.2014.01.003

